# DeepBioSim: Efficient and Versatile Methods for Microbiome Data Simulation with Minimal Statistical Assumptions

**DOI:** 10.1101/2025.08.21.670443

**Authors:** Yiyang Shen, Shri Vishalini Rajaram, Weiran Wang, Erliang Zeng

## Abstract

**Background:** The human microbiome profoundly influences health and disease. Robust computational and statistical tools for identifying causal microbe–disease links are therefore critical to uncovering the mechanistic basis of these associations. Yet benchmarking such tools remains difficult: microbiome datasets are sparse, high-dimensional, and contain complex dependencies, and no gold-standard reference set exists. Realistic simulated data with embedded ground truth are essential for fair evaluation of analytical tools. Current simulators often impose strong assumptions, require hard-to-obtain auxiliary information, or fail to scale to large, high-dimensional datasets.

**Results:** We introduce DeepBioSim, a DEEP-learning framework for BIOlogical SIMulation of microbiome data. DeepBioSim uses variational autoencoders (VAEs) to generate realistic microbiome datasets by sampling directly from the latent distribution of metagenomic or metatranscriptomic count data.

**Conclusions:** The approach is fast, accurate, and scalable, generating highly realistic synthetic microbiome datasets without extensive hyper-parameter tuning or phylogenetic input. Tests on human RNA-seq data confirm versatility of DeepBioSim, showing it can also reliably simulate single-organism omics profiles.

## 1 Introduction

The human microbiome comprises trillions of microorganisms that inhabit our bodies, forming a complex ecosystem vital to well-being. Often called the “second genome” [1], these microbial communities shape key physiological processes and are implicated in disorders such as obesity, diabetes, inflammatory bowel disease, and cancer [2, 3, 4, 5, 6]. Thoroughly characterizing these interconnected microbes is therefore a critical first step toward deepening our understanding of health and disease [7, 8, 9].

Advances in next-generation sequencing have produced large-scale count datasets across diverse modalities, including 16S/amplicon sequencing, shotgun metagenomics, and metatranscriptomics, that underpin contemporary microbiome research [10]. These datasets are inherently *compositional*, their abundances are relative to sample-specific library sizes, and exhibit extreme sparsity, zero inflation, and high dimensionality. Generating realistic synthetic microbiome data that preserve these statistical and biological properties is essential for benchmarking analytical pipelines, validating statistical methods, and supporting experimental design. However, faithfully simulating such data remains challenging due to complex feature dependencies, variable sequencing depths, and nonlinear abundance structures.

Statistical approaches for microbiome simulation often rely on rigid distributional assumptions that cannot fully capture these complexities. Dirichlet-multinomial regression models [11], gamma-hypergeometric models such as MetaSPARSim [12], and hierarchical frameworks like SparseDOSSA [13] impose fixed count distributions or hierarchical priors that may not reflect real-world variability. More advanced parametric frameworks, such as the microbiome data simulator MIDASim [14], incorporate nested distributions to better model sparsity and correlation, outperforming many deep-learning methods. Nevertheless, the parametric version of MIDASim demands extensive hyper-parameter tuning, scales poorly to datasets with thousands of taxa or genes, and is computationally prohibitive for the “narrow-rectangle” designs common in clinical microbiome studies (many features, few samples). Nonparametric methods, such as kernel density estimation (KDE) [15], struggle to scale to high-dimensional spaces and become impractical for modern microbiome datasets.

Deep generative models, in principle, can overcome these limitations by learning complex, nonlinear feature dependencies directly from data without prespecified distributional forms. Generative adversarial network (GAN) approaches, such as MB-GAN [16] and DeepMicroGen [17], have been applied to microbiome data simulation. While these methods can capture high-order relationships, they have notable drawbacks: DeepMicroGen requires auxiliary inputs such as phylogenetic or taxonomic trees, which are not always available; MB-GAN generates one sample at a time, risking overfitting and distortion of feature-level relationships; and both approaches are computationally intensive, performing optimally only on aggressively filtered datasets that may distort biological realism. Moreover, GANs aim to make synthetic samples indistinguishable from real ones, but do not explicitly model the underlying count-generating distribution, which can lead to mismatches in key statistical properties such as zero fraction, library size distributions, and inter-feature correlations.

In summary, existing simulators fall into two categories: (i) statistical methods that make strong distributional assumptions and often fail to scale, and (ii) deep-learning methods that can model complexity but require restrictive inputs, operate at the sample rather than feature level, or lack explicit count distribution modeling. However, there remains no simulator that jointly: (i) learns realistic count distributions without rigid parametric assumptions, (ii) scales to thousands of taxa/genes without aggressive feature filtering, (iii) operates without phylogenetic priors, and (iv) generalizes beyond microbiome to other omics count data.

We address this gap with **DeepBioSim**, a DEEP-learning framework for BIOlogical SIMulation of metagenomic and metatranscriptomic count data. DeepBioSim comprises three variants: DeepBioSim-VAE (standard variational autoencoder) [18], DeepBioSim-IWAE (Importance-Weighted Autoencoder) [19], and DeepBioSim-Diffusion (denoising diffusion probabilistic model) [20]. Each variant is trained directly on log-transformed count matrices to learn a high-dimensional latent representation and reconstruct realistic count profiles that preserve the sparsity, compositionality, and correlation structure of the original data. The framework scales efficiently to datasets with thousands of features, requires minimal hyper-parameter tuning, and does not depend on external phylogenetic information. Across multiple real-world microbiome datasets, all three variants generate simulated count data that closely mirror the true underlying distributions. Furthermore, when applied to a bulk RNA-seq dataset, DeepBioSim reproduced gene-expression profiles with equal fidelity, demonstrating that the framework generalizes beyond microbial communities and can accurately simulate other single-organism omics data.

## 2 Methods

### 2.1 Datasets

The datasets were stored as taxa/genes-by-sample matrices (Table 1). A `log1p` transformation was applied during training to stabilize gradients and prevent exploding back-propagation.

**Table 1.**
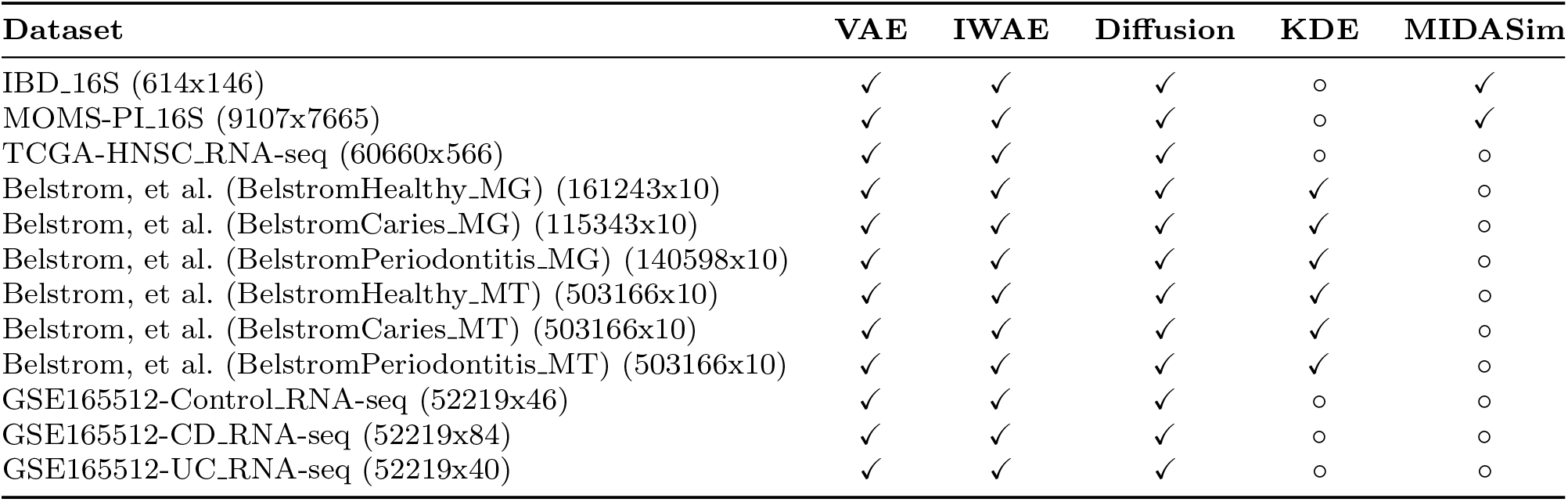
Datasets, their dimensions (features x samples), and the simulators applied in this study. For each method-dataset pair, a checkmark (✓) indicates the simulation was executed and a circle (◦) marks runs that failed because of memory or runtime constraints.

The 16S rRNA amplicon datasets for inflammatory bowel disease (IBD 16S) and the multi-omic microbiome study-pregnancy initiative (MOMS-PI 16S) were obtained from the Human Microbiome Project phase 2 (HMP2) [21]. These dataset are considered as high-dimensional because they have very large sample size. Expression profiling by high throughput sequencing for the characterization of the IL23-IL17 immune axis in IBD patients (GEO accession number: GSE165512) with 3 groups (Control, Crohn’s Disease (CD), and Ulcerative Colitis (UC)) are included as a medium-dimensional dataset considering their moderate sample size. Performance on low-dimensional data was assessed with the oral metagenomic (MG) and metatranscriptomic (MT) datasets from Belstrom et al., comprising healthy, caries, and periodontitis cohorts [22]. Finally, the TCGA head-and-neck squamous cell carcinoma RNA-seq dataset (TCGA-HNSC RNA-seq) served as a single-omics, high-dimensional test set [23]. A comprehensive summary of all datasets appears in Table 1.

### 2.2 Simulation

#### 2.2.1 Kernel Density Estimation (KDE)

KDE is a nonparametric method to estimate probability density at any point given the data. It takes the form

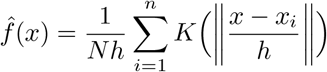

Note that K is a box function, discrete by definition, but it is common to replace it with a continuous kernel to obtain a smoother estimate. In practice, the specific kernel matters little; what truly governs estimator quality is the bandwidth *h*. When *h* → 0, the density estimate becomes jagged and overly sensitive to noise, whereas *h* → ∞ produces an oversmoothed, under-fitted curve. Bandwidth can be selected by heuristics such as Silverman’s rule of thumb or by more computationally intensive approaches like cross-validation.

Numerous improvements have been done on KDE’s computational efficiency. The naive method over *N* data points, *M* query locations in *D*-dimensional space costs O (*NMD*). Fast Fourier Transform (FFT)-based evaluators such as FFTKDE achieves better efficiency ^1^, which scales linearly with the number of points with a runtime O (2^*D*^*N* + *G* log *G*), where *G* is the number of grid cells (*G* = 2^10^ per 1-D default). It scales linearly with *N*, but explodes when increasing number of grid cells. More recent work using hash tables to estimate density has achieved good results, but the runtime can be large due to high dimensional data input and its resulting constant [15]. Those improvements fail to address the high dimensionality problem as the size of the hash table exponentially dominates the runtime.

After estimating the probability density, we used ancestral sampling to directly draw samples. Kernel density estimation (KDE) serves as a baseline, evaluated alongside MIDASim only on low-dimensional datasets (fewer than 10 samples); in higher dimensions KDE becomes impractical, the available samples cannot support an accurate density estimate, and even if they could, the runtime would be prohibitive.

#### 2.2.2 Variational Autoencoders (VAEs)

A VAE [24] models the data distribution *p*(x) by introducing a latent variable z. The encoder *q*_*φ*_(z|x) approximates the intractable posterior *p*_*φ*_(z|x), while the decoder *p*_*θ*_(x|z) with some unknown *θ* parameters to reconstructs the observations to yield 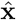. We can derive an evidence lower bound (ELBO) with the approximate posterior *q*_*φ*_(z|x)

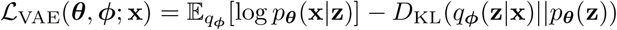

Our VAE implementation used four hidden layers with 128 units each, a 16-dimensional latent space, and is trained for 300 epochs using mini-batches of 128, the Adam optimizer, and SiLU activation. To stabilize training, log-variance values are clamped to [−5, 5] and gradient norms are clipped to a maximum of 5. We omitted the VAE for low-dimensional datasets: the latent space would be disproportionately large, risking overfitting and merely reproducing the input samples.

#### 2.2.3 Importance-weighted Autoencoders (IWAEs)

The VAE objective can be overly restrictive: when the approximate posterior deviates from the true distribution, the KL term imposes a steep penalty that limits latent expressiveness. IWAE [19] mitigates this by replacing the single-sample ELBO with a *k*-sample importance-weighted estimate of the log-likelihood, yielding a tighter lower bound and richer representations than those of the VAE:

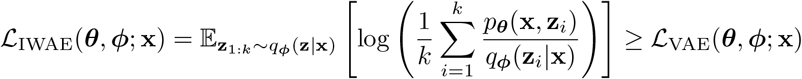

It can be further proven that the average importance weights of L_IWAE_ are unbiased estimators of *p*(x) trivially using Jensen’s inequality, and that it approaches log *p*(x) as *k* goes to infinity.

The implementation hyperparameters are identical to those of the vanilla VAE. The only change for IWAE is that we used 20 weights (*k* = 20) to improve generation quality.

#### 2.2.4 Denoising Diffusion Probabilistic Models (Diffusion Models)

Denoising diffusion probabilistic models [20] also rely on latent variables *z*, much like VAEs, but yet they differ fundamentally from VAEs by using a Markov-chain framework. The forward (noise-adding) process

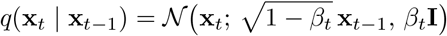

gradually injects Gaussian noise, pushing the data distribution *p*(x_0_) toward the standard normal N (0, I). Unlike VAEs, the latent variable is sampled from this noisy trajectory rather than from a learned prior.

Because the forward process admits a closed-form solution, we can sample x_*t*_ directly from

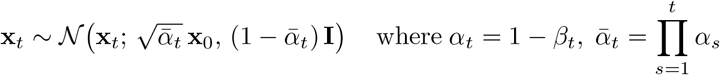

or equivalently,

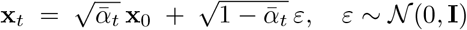

The reverse (denoising) process is modeled as a chain of Gaussians parameterized by *θ*:

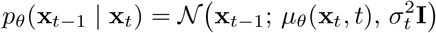

where *µ*_*θ*_(x_*t*_, *t*) is computed via a network *ε*_*θ*_(x_*t*_, *t*). The network is trained by minimizing the variational upper bound on the negative log-likelihood, which reduces to:

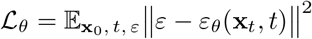

Our implementation uses 3,000 diffusion steps, 128 hidden units, and a linear *β*_*t*_ schedule.

### 2.3 Simulation Evaluation

#### 2.3.1 Runtime

Efficiency was measured by recording the wall-clock runtime (in seconds) on CPU and GPU. MIDASim cannot leverage GPU acceleration, whereas DeepBioSim runs natively on CUDA- and MPS-enabled hardware.

#### 2.3.2 Visualization

We evaluate simulation quality with unsupervised visualizations including PCA, t-SNE, and UMAP, and with violin plots of alpha-diversity metrics (richness and Shannon index).

#### 2.3.3 Sample-level Properties

For each simulation method, we quantified fidelity by computing Pearson correlations between diversity metrics from the real and simulated samples. We compared alpha-diversity (richness, Shannon index) across samples and beta-diversity (Bray–Curtis, Jaccard) across sample pairs; coefficients values closer to 1 indicate stronger concordance.

#### 2.3.4 Assessing Fidelity of Simulated Data through Classification Performance

To evaluate the generation fidelity of each model, we trained a single-layer MLP (ReLU activation, Adam optimizer, 2,000 iterations, batch size = 16) and a linear classifier to distinguish among the three conditions in the GSE165512 RNA-seq dataset: control, ulcerative colitis (UC), and Crohn’s disease (CD). Because all groups shared the same gene features, we first reduced dimensionality to 40 components using kernel PCA. Each classifier was trained 100 times on either real or simulated data and evaluated on a held-out test set comprising 20% of the original samples [25]. We assessed generative quality by computing the inter-entropy and intra-entropy of predictions for the simulated data. Inter-entropy measures the diversity of generated samples based on the predicted label distribution *p*(*y*); higher values indicate greater diversity. Intra-entropy measures prediction uncertainty; lower values indicate higher classifier confidence.

## 3 Results

### 3.1 Runtime

We benchmarked the three DeepBioSim variants (DeepBioSim-VAE, DeepBioSim-IWAE, and DeepBioSim-Diffusion) against the KDE and MIDASim simulators. Runtime benchmarking was conducted on 12 omics datasets: eight microbiome sets and four human RNA-seq sets. The microbiome data span 16S rRNA amplicon sequencing (16S), whole-metagenome shotgun sequencing (MG), and metatranscriptomics (MT); dataset characteristics and associated diseases are listed in Table 1. As summarized in Table 2, all three DeepBioSim variants are computationally efficient, with DeepBioSim-Diffusion proving fastest on higher-dimensional data (IBD 16S and MOMS-PI 16S). Compared with the DeepBioSim variants, KDE is fastest on low-dimensional data (BelstromHealthy MG, BelstromCaries MG, BelstromPeriodontitis MG, BelstromHealthy MT, BelstromCaries MT, and BelstromPeriodontitis MT), whereas MIDASim remains the slowest regardless of dimensionality. Four high-dimensional RNA-seq datasets (TCGA-HNSC RNA-seq, GSE165512-Control RNA-seq, GSE165512-CD RNA-seq, and GSE165512-UC RNA-seq) were also benchmarked and similar results were obtained, demonstrating the versatility of our framework. The DeepBioSim variants outperformed MIDASim and KDE on these datasets, with DeepBioSim-Diffusion being the fastest. The results demonstrate that DeepBioSim is not only fast but also scales well to high-dimensional datasets, making it a practical choice for microbiome data simulation.

**Table 2.** Running times (in seconds) for each method across different datasets. The most efficient method for each dataset is highlighted in bold. The dashes (—) indicate that the method was not applicable or not executable on that dataset (see Table 1 for details).

| Dataset | DeepBioSim-VAE | DeepBioSim-IWAE | DeepBioSim-Diffusion | KDE | MIDASim |
| --- | --- | --- | --- | --- | --- |
| IBD_16S | 10.5775 | 12.6244 | 10.0078 | — | **8.0840** |
| MOMS-PI_16S | **155.5329** | 353.6716 | 312.9778 | — | 10959.0270 |
| Belstrom, et al. (BelstromHealthy_MG) | 1952.8125 | 2628.1077 | 804.8901 | **0.0405** | — |
| Belstrom, et al. (BelstromCaries_MG) | 1399.8902 | 1873.2686 | 568.2599 | **0.0427** | — |
| Belstrom, et al. (BelstromPeriodontitis_MG) | 1701.9459 | 2280.5595 | 724.4212 | **0.0759** | — |
| Belstrom, et al. (BelstromHealthy_MT) | 6430.0490 | 8262.6575 | 2525.0858 | **0.0872** | — |
| Belstrom, et al. (BelstromCaries_MT) | 5984.7729 | 8239.0809 | 2481.6411 | **0.0938** | — |
| Belstrom, et al. (BelstromPeriodontitis_MT) | 6163.5157 | 8242.2774 | 2483.6330 | **0.0947** | — |
| TCGA-HNSC.RNA-seq | 766.1962 | 1069.3793 | **388.9872** | — | — |
| GSE165512-Control.RNA-seq | 654.4115 | 882.7418 | **272.0061** | — | — |
| GSE165512-CD.RNA-seq | 648.3288 | 863.7208 | **268.8331** | — | — |
| GSE165512-UC.RNA-seq | 663.0517 | 883.4175 | **267.2024** | — | — |

### 3.2 Gene Level/Taxon-level Comparison Between Simulated and Original Data

Comparison with the original dataset shows that DeepBioSim produces simulated microbiome count data that closely mirror the real observations. The simulations preserve complex inter-taxa/inter-gene relationships, confirming the method’s ability to generate realistic microbiome data.

Across diverse high-dimensional microbiome datasets, including the MOMS-PI 16S cohort (MOMS-PI 16S, Figure 1) and an IBD 16S cohort (IBD 16S, Figure 2), DeepBioSim-IWAE generated the most accurate simulations, closely matching the original taxon abundances and gene-expression levels. DeepBioSim-VAE also performed well, though its simulated gene distributions were slightly less dispersed than in the real data. DeepBioSim-Diffusion and MIDASim yielded the least representative profiles. These performance rankings remained consistent across all three dimensionality-reduction visualisations (Figures 1 and 2), including principal component analysis (PCA), t-distributed stochastic neighbor embedding (t-SNE), and uniform manifold approximation and projection (UMAP).

**Fig. 1.**
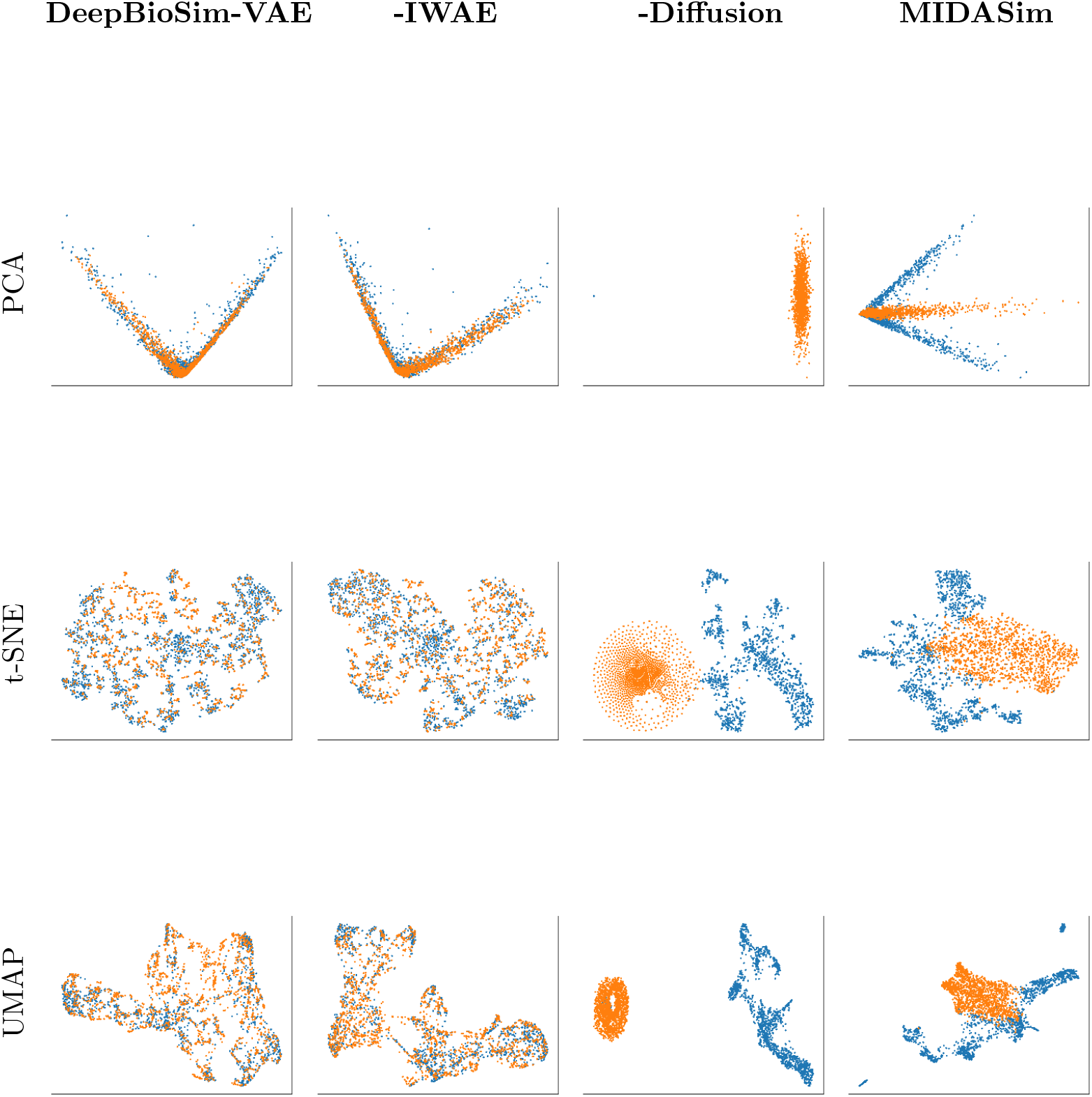
Taxon-level concordance between the original microbiome dataset MOMS-PI 16S and four simulated counterparts. Blue points represent taxa in the original data; orange points represent taxa in the simulated data. Each dimensionality-reduction technique visualises how closely the simulated taxon-abundance profiles resemble the originals for each modelling approach.

**Fig. 2.**
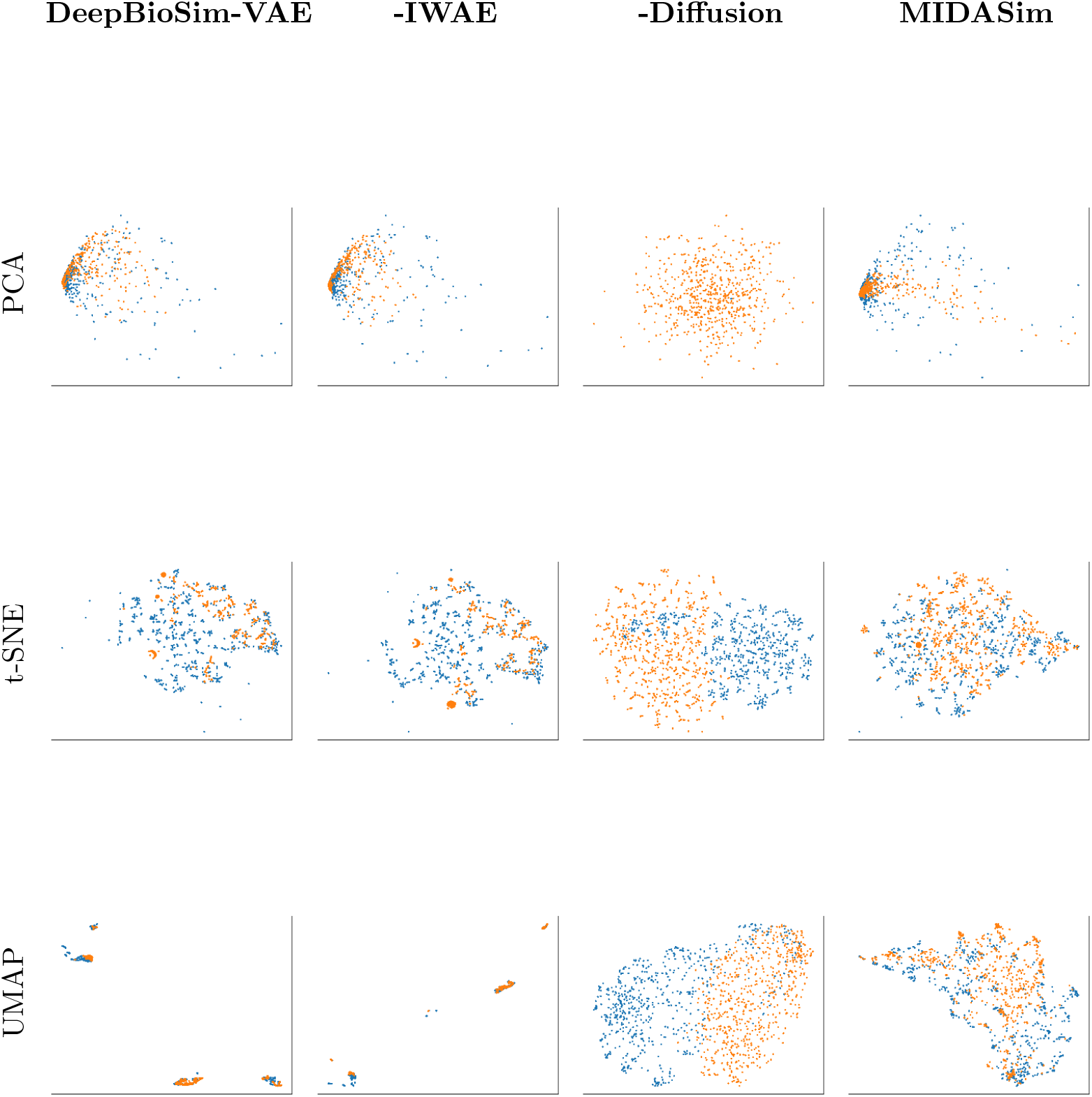
Taxon-level concordance between the original microbiome dataset IBD 16S and four simulated counterparts. Blue points represent taxa in the original data; orange points represent taxa in the simulated data. Each dimensionality-reduction technique visualizes how closely the simulated taxon-abundance profiles resemble the originals for each modelling approach.

Consistent performance patterns appeared in the high-dimensional TCGA-HNSC RNA-seq dataset (TCGA-HNSC RNA-seq, Figure A5) and GSE165512 RNA-seq cohorts (Control, CD, and UC, Figures A6, A7, and A8). Memory demand of MIDASim grows exponentially with dataset size, so it exceeded our available resources (128-core 512G RAM on high-performance computers (HPC)) and could not be evaluated on the high-dimensional (large sample number) datasets.

Our methods thrive partly because of its ability to be run in rectangular datasets common in clinical settings (few samples but many taxa). In the low-dimensional MG and MT datasets from Belstrom et al. [22], all three of our proposed methods (DeepBioSim-VAE, DeepBioSim-IWAE, and DeepBioSim-Diffusion) have great performance and is on par with kernel density estimation (KDE) baseline method, which can be used in a low-dimensional setting (Figures 3, 4, A1, A2, A3, A4). MIDASim could not be evaluated, as its memory requirements were too large and exceeded our available resources (128-core 512G RAM on high-performance computers (HPC)).

**Fig. 3.**
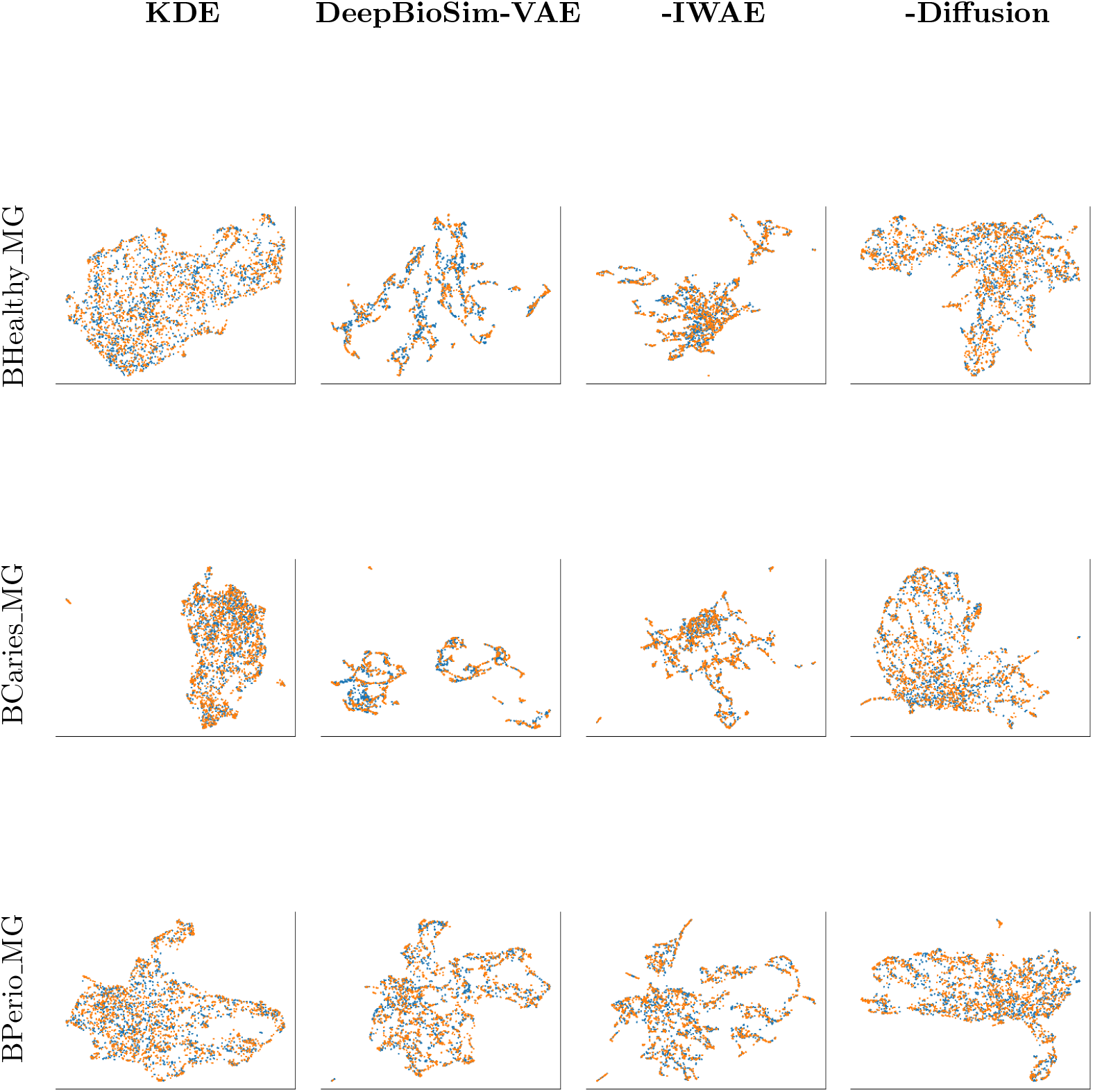
Gene-level concordance between the original BelstromHealthy MG, Bel-stromCaries MG, and BelstromPeriodontitis MG microbiome datasets and their two simulated counterparts, using UMAP visualization. Blue points represent genes in the original data; orange points represent genes in the simulated data.

**Fig. 4.**
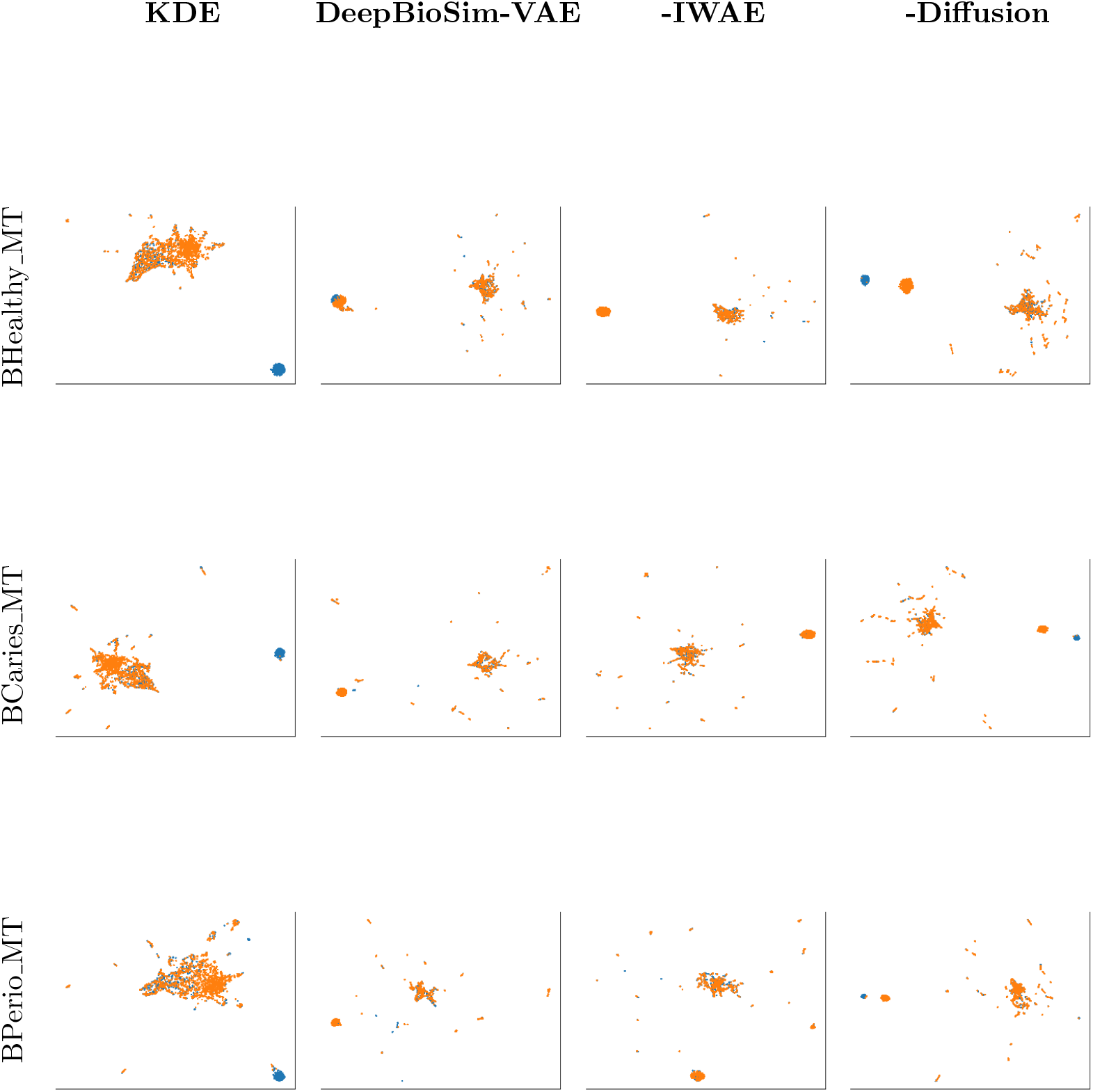
Gene-level concordance between the original BelstromHealthy MT, Bel-stromCaries MT, and BelstromPeriodontitis MT microbiome datasets and their two simulated counterparts, using UMAP visualization. Blue points represent genes in the original data; orange points represent genes in the simulated data.

### 3.3 Sample Level Properties Evaluation

While DeepBioSim targets taxon-level/gene-level patterns, the resulting synthetic datasets also accurately preserve the sample-level characteristics of the original data. Benchmarks comparing alpha-diversity (richness and Shannon indices) and beta-diversity (Bray-Curtis and Jaccard dissimilarity) between real and simulated samples show close concordance, confirming that DeepBioSim accurately reproduces both richness and evenness, and thus generates microbiome data that are realistic at community and sample scales.

In the high-dimensional 16S datasets (IBD 16S and MOMS-PI 16S), DeepBioSim-IWAE and MIDASim preserved sample-level diversity most reliably, closely matching the observed richness and Shannon entropy (Figures 5, 6). DeepBioSim-VAE followed, though its simulations showed slightly greater dispersion in these metrics. DeepBioSim-Diffusion failed to capture the diversity profile (Figure B9, B10).

**Fig. 5.**
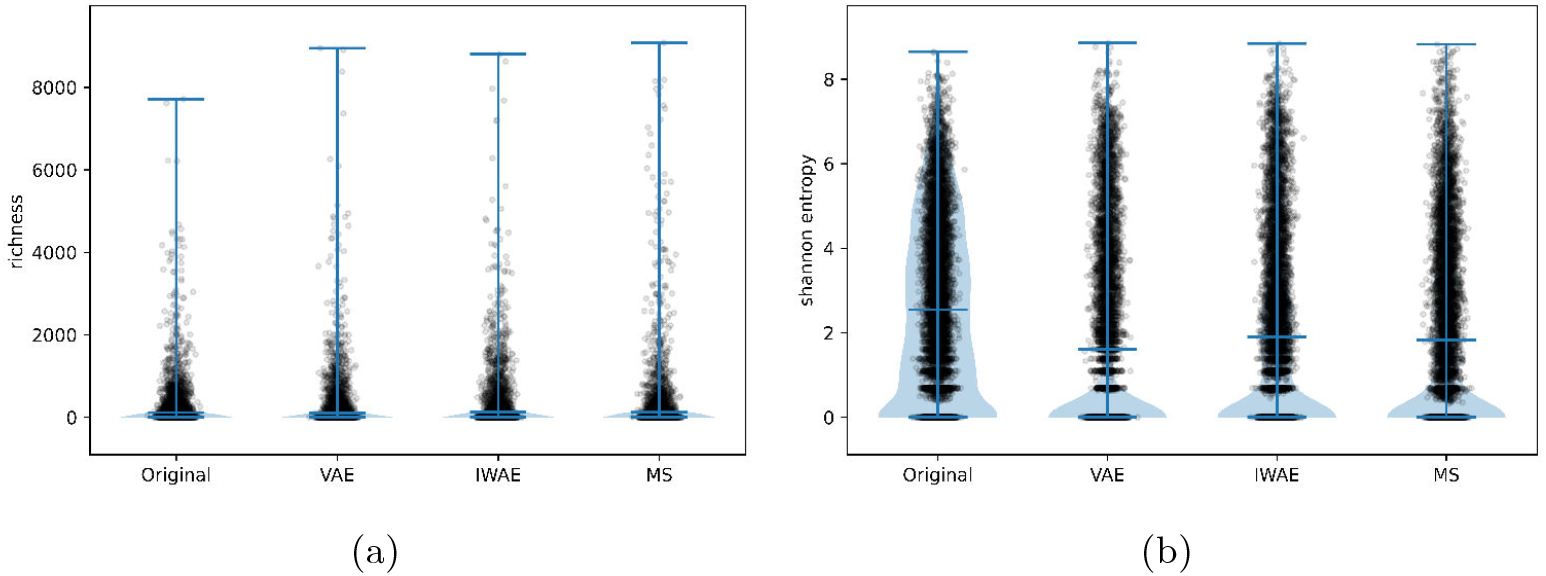
Sample-level alpha-diversity concordance between the original MOMS-PI 16S dataset and four simulated datasets. Panels (a) and (b) plot richness and Shannon entropy, respectively, after omitting the DeepBioSim-Diffusion model to improve visual clarity. Each point represents a single sample. MS refers to method MIDASim.

**Fig. 6.**
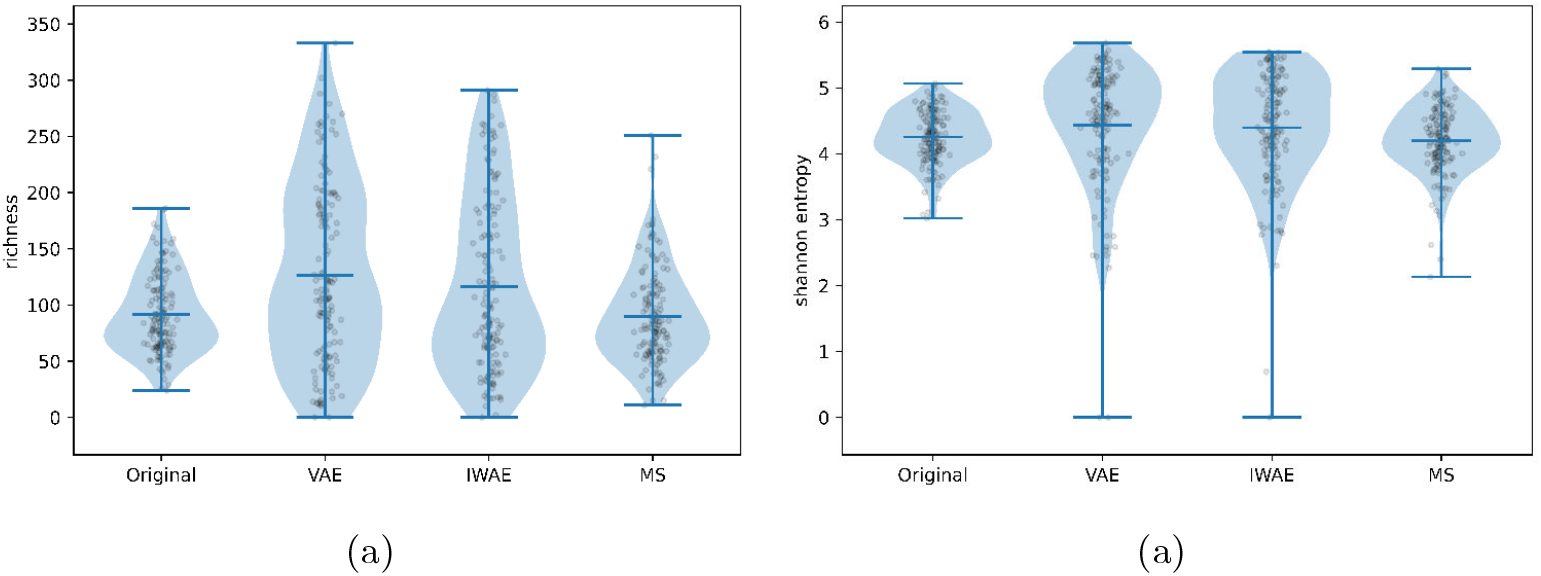
Sample-level alpha-diversity concordance between the original MOMS-PI 16S dataset and four simulated datasets. Panels (a) and (b) plot richness and Shannon entropy, respectively, after omitting the DeepBioSim-Diffusion model to improve visual clarity. Each point represents a single sample. MS refers to method MIDASim.

We quantified these findings by computing Pearson correlation coefficients between real and simulated samples for alpha-diversity (richness and Shannon) and beta-diversity (Bray-Curtis and Jaccard). DeepBioSim-IWAE consistently achieved the highest correlations, confirming its superior fidelity; DeepBioSim-VAE ranked next, whereas MIDASim lagged well behind (Tables 3, 4). DeepBioSim-Diffusion was excluded from these metrics for MOMS-PI 16S and IBD 16S due to its poor performance in capturing gene-and sample-level properties, which led to overflow errors in metric calculations (See Figures 1, 2, B9, B10).

**Table 3.** Pearson Correlation Coefficient of Diversity Index between Simulated MOMS-PI 16S Datasets and Original Dataset.

| Metric | DeepBioSim-VAE | DeepBioSim-IWAE | MIDASim |
| --- | --- | --- | --- |
| Shannon | 0.7613 | **0.8225** | 0.7190 |
| Richness | **0.9698** | 0.9695 | 0.7511 |
| Bray-Curtis | -0.0531 | **-0.0221** | -0.0564 |
| Jaccard | -0.0577 | **-0.0128** | -0.0502 |

**Table 4.** Pearson Correlation Coefficient of Diversity Index between Simulated IBD 16S Datasets and Original Dataset.

| Metric | DeepBioSim-VAE | DeepBioSim-IWAE | MIDASim |
| --- | --- | --- | --- |
| Shannon | **0.7928** | 0.5934 | 0.4661 |
| Richness | **0.9156** | 0.9093 | 0.8026 |
| Bray-Curtis | 0.4941 | **0.5581** | 0.0586 |
| Jaccard | 0.6812 | **0.7241** | 0.3990 |

Consistent performance patterns appeared in the additional high-dimensional TCGA-HNSC RNA-seq dataset (Table B7, Figure B13) and GSE165512 RNA-seq cohorts under different conditions (Tables B8, B9, B10, Figure B14). MIDASim was again excluded from evaluation because its substantial memory requirements made it infeasible to run on large datasets.

In the low-dimensional metagenomic and metatranscriptomic datasets from Belstrom et al., three DeepBioSim methods (VAE, IWAE, and Diffusion) and KDE perform similarly in terms of rank-based correlation coefficients of alpha and beta diversities, showing all methods are capable of capturing original data’s diversities. DeepBioSim-Diffusion surpasses KDE only on the Jaccard index, a result likely attributable to the limited number of features in these datasets (Tables B1, B2, B3, B4, B5, B6). However, violin plots, which show the absolute alpha diversity comparisons, demonstrated that DeepBioSim-IWAE performs the best out of all methods, followed closely by DeepBioSim-VAE (Figures B11 B12).

### 3.4 Assessing Fidelity of Simulated Data through Classification Performance

The heterogeneity of our benchmarking datasets, including several multi-class cohorts, allowed us to assess simulation quality both globally and within specific sample groups (Tables 5, 6) for the GSE165512 RNA-seq cohort. Multilayer perceptron (MLP) and linear classifiers trained on dataset generated by DeepBioSim-VAE and DeepBioSim-IWAE consistently outperformed those trained on DeepBioSim-Diffusion outputs (Tables 5, 6), demonstrating better preservation of group-specific signals. We did not perform this analysis on the Belstrom datasets because each condition has only 10 samples but more than 100,000 taxa; such small sample size makes dimentionality-reduction and classification unstable and thus uninformative.

**Table 5.** MLP Prediction Performance for Datasets Generated by Different Methods (95% CI). An upward arrow (↑) indicates that a larger value reflects better performance, while a downward arrow (↓) indicates

| Method | Inter-entropy $H(y) \uparrow$ | Intra-entropy $H(y\vert x) \downarrow$ |
| --- | --- | --- |
| DeepBioSim-VAE | **1.4914 (1.4267-1.5380)** | **0.1570 (0.0546-0.3138)** |
| DeepBioSim-IWAE | 1.4840 (1.3934-1.5411) | 0.3428 (0.2148-0.5335) |
| DeepBioSim-Diffusion | 1.3849 (1.1669-1.5183) | 0.4811 (0.3388-0.6621) |

**Table 6.** Linear Prediction Performance for Datasets Generated by Different Methods (95% CI). An upward arrow (↑) indicates that a larger value reflects better performance, while a downward arrow (↓) indicates that a smaller value reflects better performance.

| Method | Inter-entropy $H(y) \uparrow$ | Intra-entropy $H(y\vert x) \downarrow$ |
| --- | --- | --- |
| DeepBioSim-VAE | 1.4976 (1.4497-1.5417) | **0.7140 (0.6281-0.8232)** |
| DeepBioSim-IWAE | **1.5021 (1.4427-1.5503)** | 0.8755 (0.7774-0.9965) |
| DeepBioSim-Diffusion | 1.4832 (1.4039-1.5496) | 1.0983 (1.0134-1.2060) |

## 4 Discussion

Our results demonstrate that DeepBioSim can reliably reproduce both taxon-level/gene-level patterns and sample-level properties across diverse microbiome and RNA-seq datasets. The simulated profiles preserve key ecological structures such as alpha diversity, beta diversity, and label entropy, metrics essential for maintaining biological interpretability and enabling downstream analyses including classification, clustering, and diversity-based ecological inference. This fidelity allows simulated data to serve as a valuable resource for benchmarking and validating analytical algorithms without compromising interpretability. Among the three variants, DeepBioSim-IWAE delivers the most accurate simulations in medium- and high-dimensional settings, whereas DeepBioSim-VAE comes a close second and completes runs significantly faster. In the high-dimensional MOMS-PI 16S dataset, DeepBioSim-IWAE achieved the highest fidelity, with Pearson correlations to real data of *r* = 0.8225 for Shannon diversity and *r* = 0.9695 for richness, and the smallest divergence in beta-diversity (Bray–Curtis *r* = −0.0221, Jaccard *r* = −0.0128) among all tested methods (Table 3). In the IBD 16S dataset, DeepBioSim-VAE yielded the highest alpha-diversity correlations (Shannon *r* = 0.7928, richness *r* = 0.9156), while DeepBioSim-IWAE slightly outperformed it on beta-diversity (Bray–Curtis *r* = 0.5581, Jaccard *r* = 0.7241) (Table 4). Across datasets, DeepBioSim variants consistently maintained diversity correlations above 0.75 for at least one alpha and one beta metric, demonstrating robust preservation of community structure.

Architectural differences explain the observed performance hierarchy. DeepBioSim-IWAE leverages a tighter variational bound [19], enabling more accurate latent representations in medium- and high-dimensional regimes where complex inter-feature dependencies are prevalent. This advantage is evident in the MOMS-PI 16S set, where beta-diversity correlations of IWAE exceeded VAE by 3–4 percentage points despite slightly lower richness correlation. DeepBioSim-VAE, while marginally less accurate in some high-dimensional settings, converged substantially faster, like, 155.53 s vs. 353.67 s for IWAE on MOMS-PI 16S, making it attractive when computational efficiency is a priority. DeepBioSim-Diffusion, consistent with findings in other omics contexts [20], retained strong realism in low-dimensional Belstrom MG/MT datasets (UMAP/PCA visual concordance comparable to KDE) but degraded in high-dimensional data, often failing diversity preservation to the point of metric overflow.

Baseline comparisons reinforce these trade-offs. MIDASim produced plausible simulations, with some diversity correlations rivaling DeepBioSim (richness *r* = 0.7511 on MOMS-PI 16S), but runtimes were prohibitive, 10,959 s for MOMS-PI 16S versus 155–313 s for DeepBioSim variants, and memory requirements exceeded 512 GB for large datasets. KDE, although fastest in low-dimensional MG/MT datasets (0.04–0.09 s), failed to capture higher-order correlation structures and was infeasible in high-dimensional regimes. Other published methods were excluded for specific reasons: DeepMicroGen targets imputation rather than simulation [17], and MB-GAN [16] requires *>*100,000 iterations to converge, depends on deprecated libraries, and generates samples one at a time, preventing learning of population-wide structure.

DeepBioSim also preserved group-discriminative signals, as shown by classification experiments in the multi-class GSE165512 RNA-seq dataset. For MLP classifiers, DeepBioSim-VAE achieved the highest inter-entropy (*H*(*y*) = 1.4914, 95% CI: 1.4267–1.5380) and lowest intra-entropy (*H*(*y*|*x*) = 0.1570, 95% CI: 0.0546–0.3138), followed by IWAE. Similar trends held for linear classifiers, where IWAE yielded the highest inter-entropy (*H*(*y*) = 1.5021) but VAE maintained the lowest intra-entropy (*H*(*y*|*x*) = 0.7140), indicating better within-class compactness.

Despite these strengths, several limitations remain. The current implementation does not encode phylogenetic or taxonomic information, which could be critical for preserving evolutionary signal in simulated co-occurrence patterns. Future versions could integrate tree-based distances into the latent representation or add topology-aware regularization to the loss function. In datasets where samples vastly outnumber features, the model may suffer from reduced fidelity due to the curse of dimensionality; dimensionality-reduction strategies such as principal-component analysis or sparse factorization should be systematically explored. While the present framework operates on the full feature matrix, a sample-wise generation mode is feasible: compressing taxa counts via singular-value decomposition, training in reduced space, and reconstructing full matrices losslessly when the reduced dimension does not exceed min(*n, p*) [26]. The framework does not enforce one-to-one feature matching between simulated and original matrices; post-generation alignment methods such as Gale–Shapley matching [27] or conditional VAEs [28] could better preserve fine-scale feature correspondence.

In the broader context of simulation for biological data, our findings align with trends in deep generative modeling where increased flexibility in latent space modeling improves fidelity but may increase computational cost [29, 30, 31]. The consistent performance of DeepBioSim across both microbiome and RNA-seq datasets underscores its versatility, and its independence from phylogenetic priors enhances applicability to less-characterized microbial communities and single-organism transcriptomes. By making realistic simulation more accessible, DeepBioSim can support reproducible benchmarking of statistical and machine-learning workflows, power analysis for experimental design, and accelerated development of analysis tools across diverse omics domains.

## 5 Conclusions

We introduce DeepBioSim, which leverages variational autoencoders (VAEs) to generate realistic microbiome datasets by sampling directly from the latent distribution of metagenomic or metatranscriptomic count data. The approach is fast, accurate, scalable, and easy to use. Tests on human RNA-seq data confirm versatility of DeepBioSim, showing it can also reliably simulate single-organism omics profiles. By enabling the simulation of large-scale datasets, it holds significant potential for improving the benchmarking and development of computational tools in microbiome research and beyond.

## 6 List of Abbreviations

DeepBioSim: a DEEP-learning framework for BIOlogical SIMulation of microbiome data
VAE: Variational Autoencoder
IWAE: Importance Weighted Autoencoder
KDE: Kernel Density Estimation
MLP: Multi-Layer Perceptron
GAN: Generative Adversarial Network
IBD: Inflammatory Bowel Disease
MOMS-PI: Multi-Omic Microbiome Study-Pregnancy Initiative
HMP2: Human Microbiome Project 2
CD: Crohn’s Disease
UC: Ulcerative Colitis
MG: Metagenomic
MT: Metatranscriptomic
TCGA-HNSC: The Cancer Genome Atlas Head and Neck Squamous Cell Carcinoma
FFT: Fast Fourier Transform
SiLU: Sigmoid Linear Unit
DDPM: Denoising Diffusion Probabilistic Models
PCA: Principal Component Analysis
t-SNE: t-distributed Stochastic Neighbor Embedding
UMAP: Uniform Manifold Approximation and Projection

## 7 Declarations

### 7.1 Ethics approval and consent to participate

Not applicable.

### 7.2 Consent for publication

Not applicable.

### 7.3 Availability of Data and Material

Data used in this study, their corresponding processing pipeline, and code for the simulation methods are available at https://github.com/biocoms/DeepBioSim.

### 7.6 Authors’ Contributions

YS: Methodology, Software, Analysis, Writing

SVR: Methodology, Writing

WW: Methodology, Writing

EZ: Conception, Supervision, Methodology, Writing

## 7.7 Acknowledgements

We acknowledge the University of Iowa ITS-Research Services for access to the Argon high-performance computing system, which supported several of our experiments, and Mengyu He, creator of MIDASim, for generously sharing analysis code and clarifying methodological details.

## Appendix A Additional Visualizations

### A.1 Visualizations for datasets from Belstrom et al

**Fig. A1.**
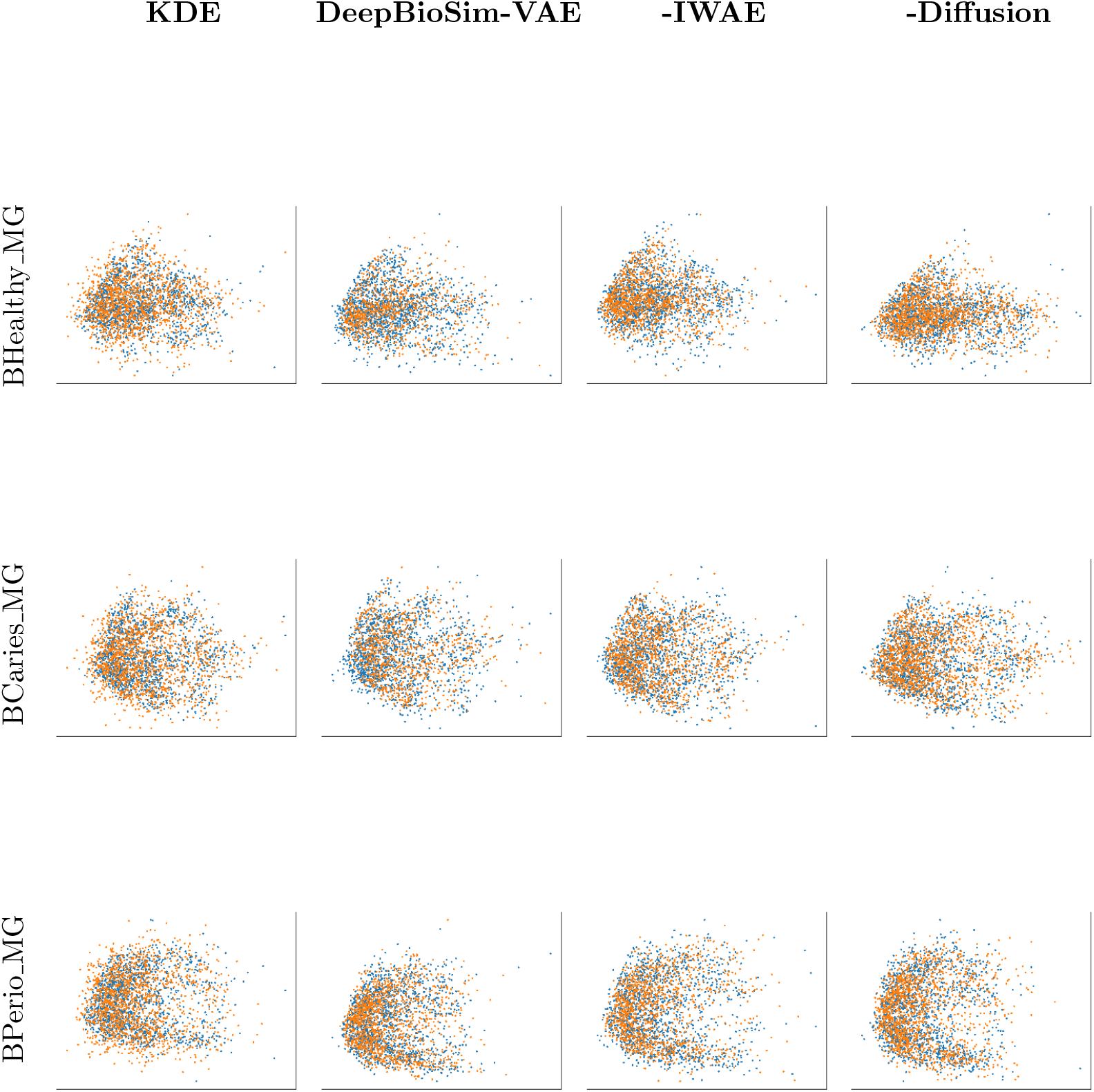
Gene-level concordance between the original BelstromHealthy MG, Bel-stromCaries_MG, and BelstromPeriodontitis MG microbiome datasets and their two simulated counterparts, using PCA visualization. Blue points represent genes in the original data; orange points represent genes in the simulated data.

**Fig. A2.**
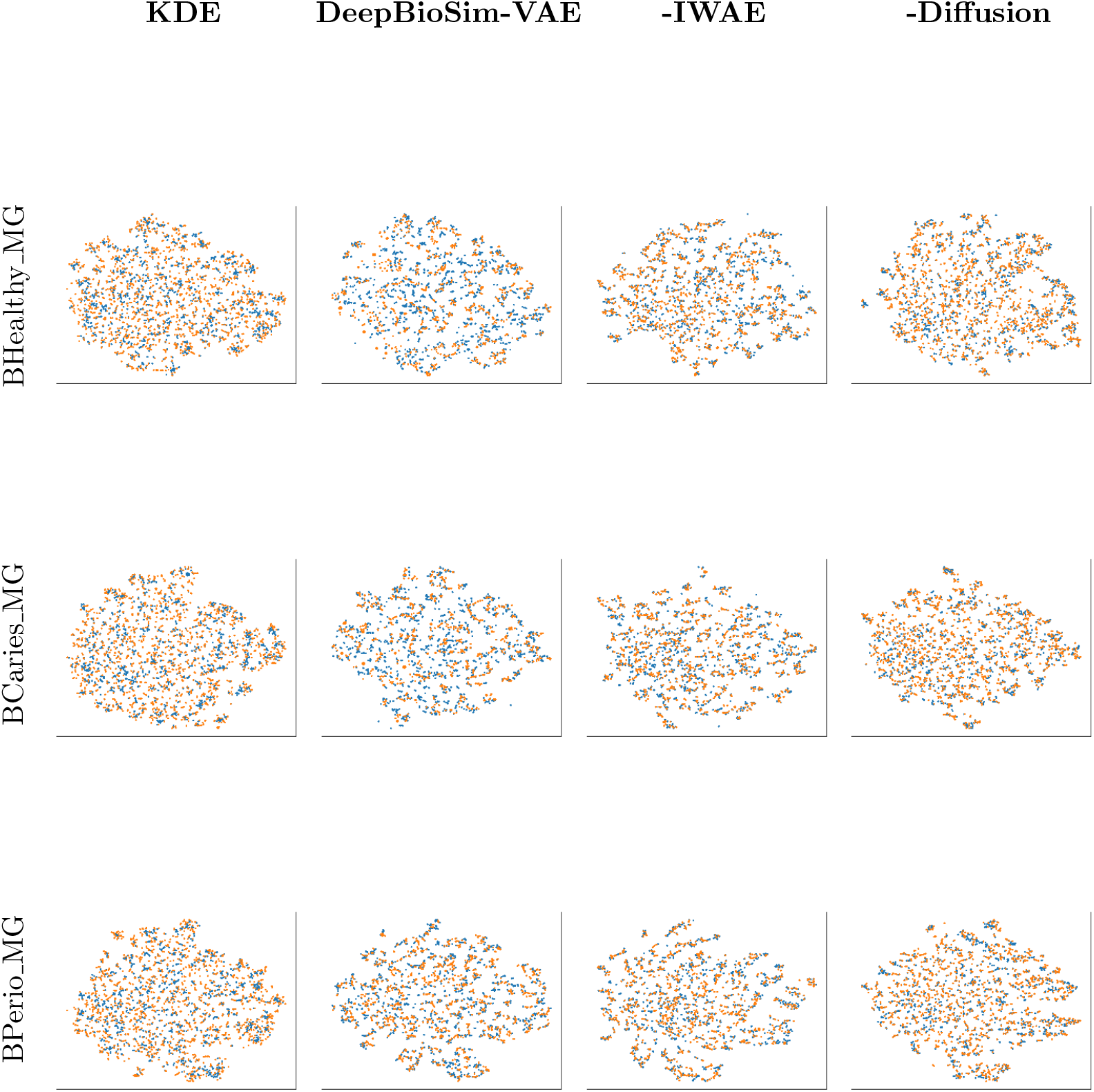
Gene-level concordance between the original BelstromHealthy MG, Bel-stromCaries_MG, and BelstromPeriodontitis MG microbiome datasets and their two simulated counterparts, using t-SNE visualization. Blue points represent genes in the original data; orange points represent genes in the simulated data.

**Fig. A3.**
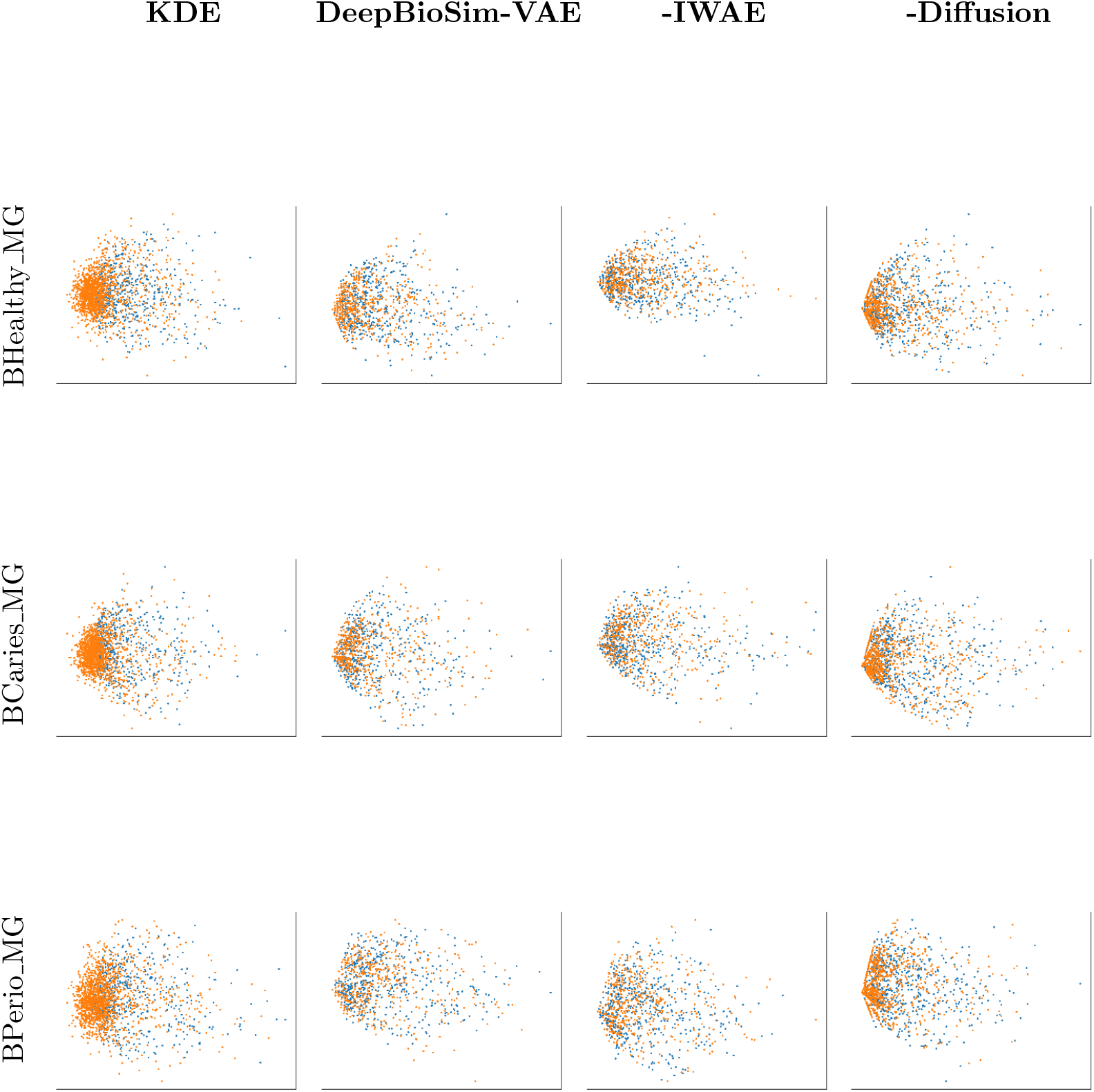
Gene-level concordance between the original BelstromHealthy MT, Bel-stromCaries MT, and BelstromPeriodontitis MT microbiome datasets and their two simulated counterparts, using PCA visualization. Blue points represent genes in the original data; orange points represent genes in the simulated data.

**Fig. A4.**
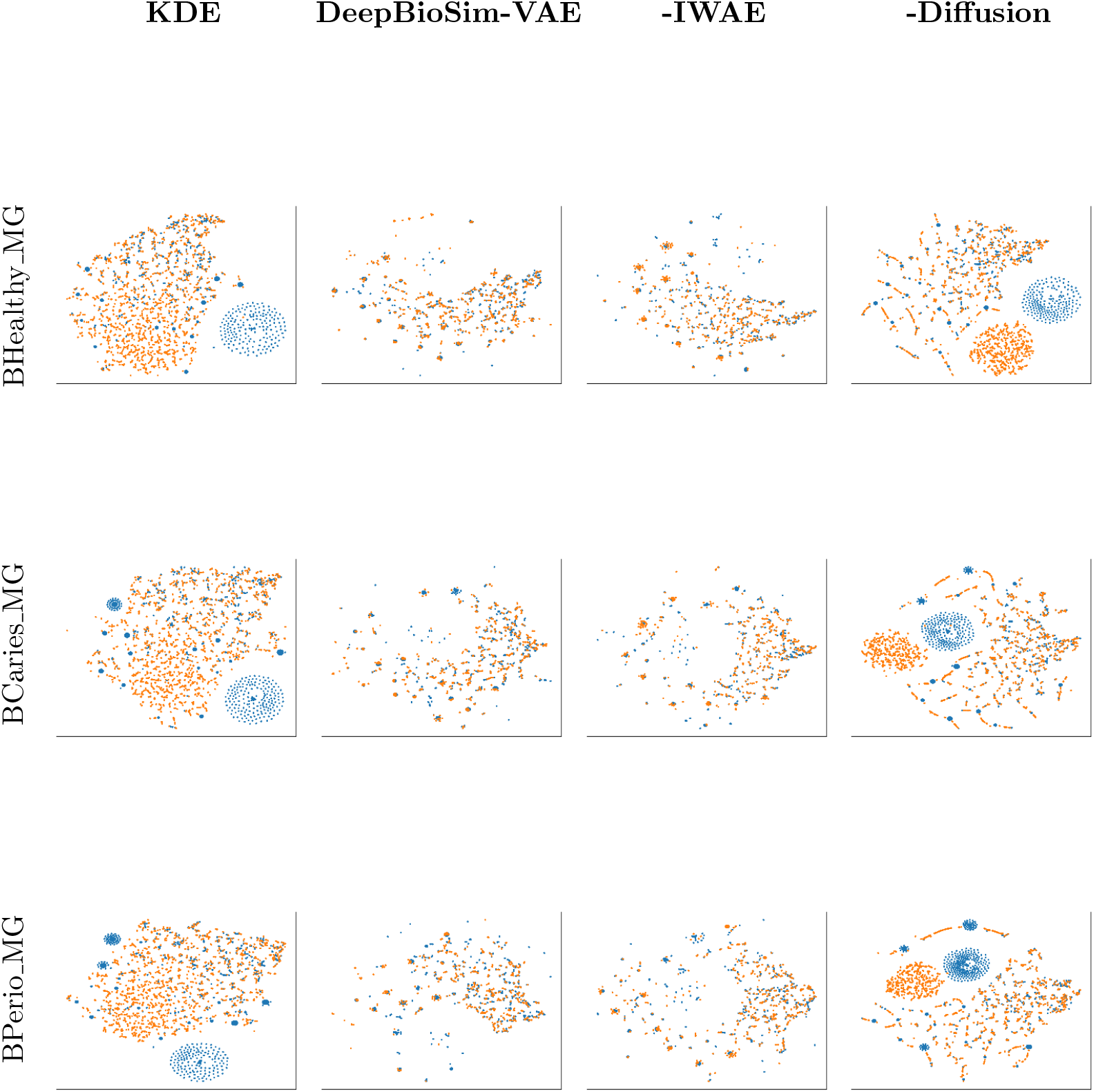
Gene-level concordance between the original BelstromHealthy MT, Bel- stromCaries MT, and BelstromPeriodontitis MT microbiome datasets and their two simulated counterparts, using t-SNE visualization. Blue points represent genes in the original data; orange points represent genes in the simulated data.

### A.2 Visualizations for TCGA-HNSC RNA-seq dataset

**Fig. A5.**
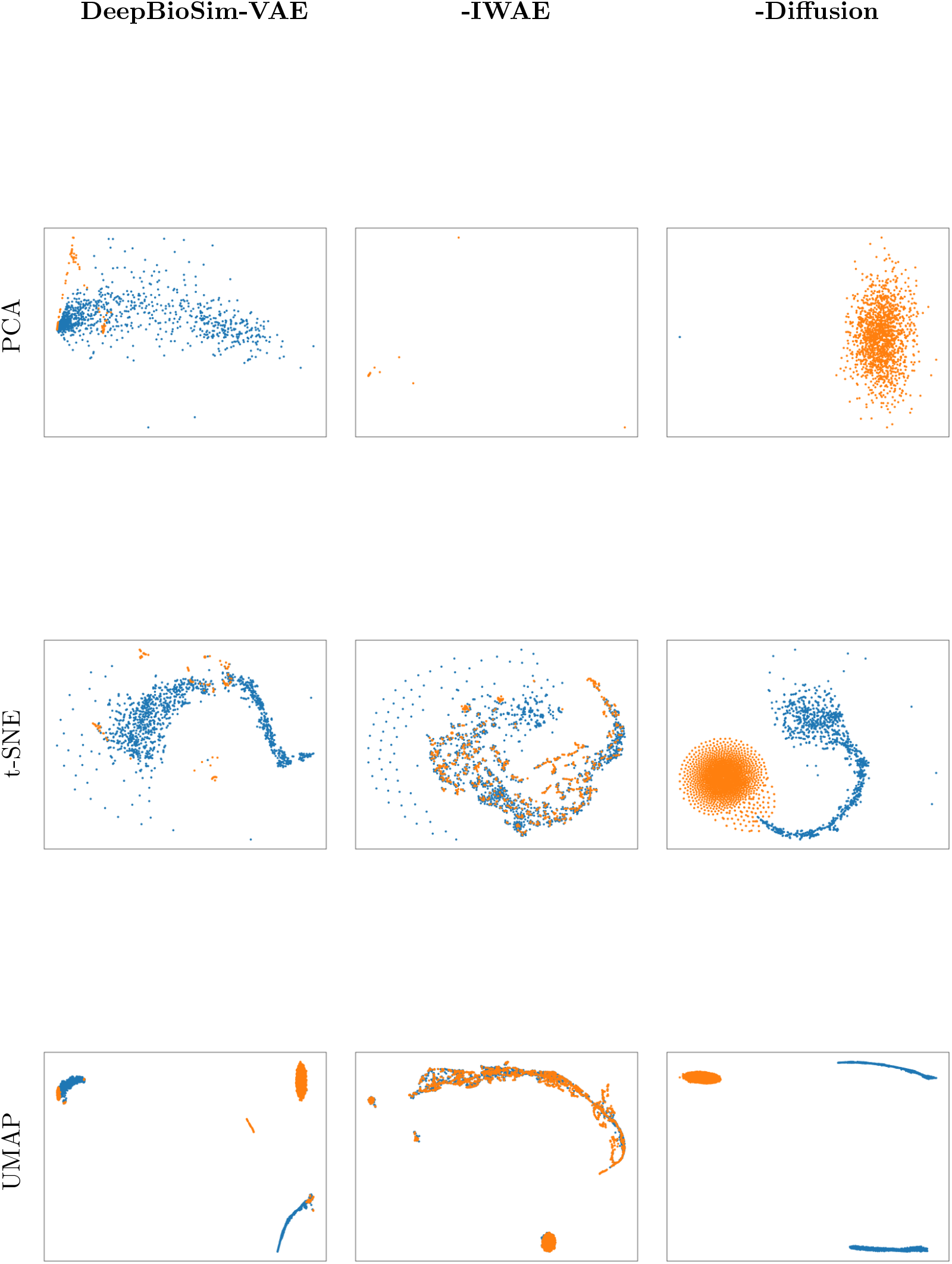
Gene-level concordance between the original microbiome dataset TCGA- HNSC RNA-seq and four simulated counterparts. Blue points represent genes in the original data; orange points represent genes in the simulated data.

### A.3 Visualizations for datasets from GSE165512

**Fig. A6.**
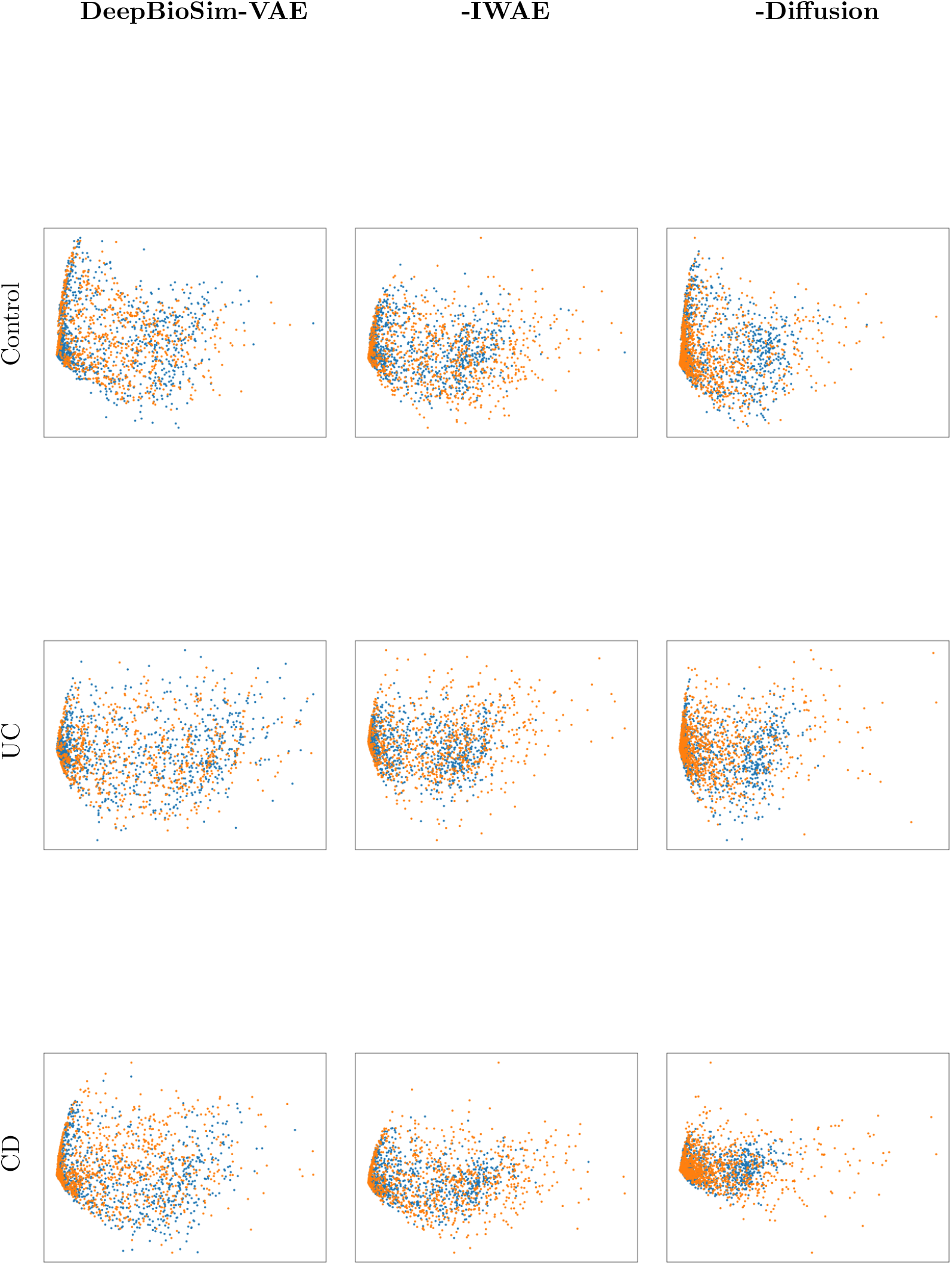
Gene-level concordance between the original human RNA-seq datasets from GSE165512 with different conditions and their three simulated counterparts, using PCA visualization. Blue points represent genes in the original data; orange points represent genes in the simulated data.

**Fig. A7.**
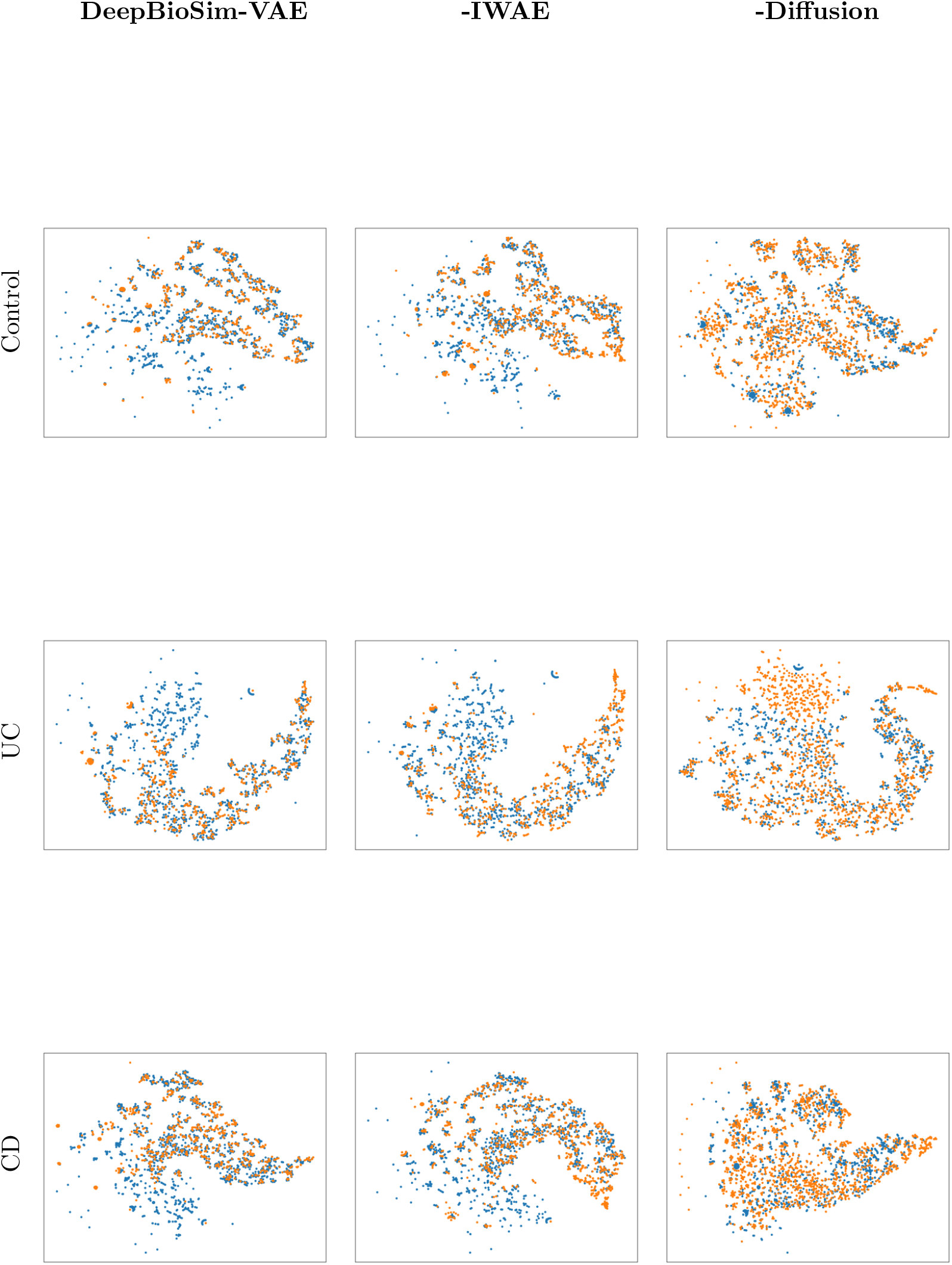
Gene-level concordance between the original human RNA-seq datasets from GSE165512 with different conditions and their three simulated counterparts, using t-SNE visualization. Blue points represent genes in the original data; orange points represent genes in the simulated data.

**Fig. A8.**
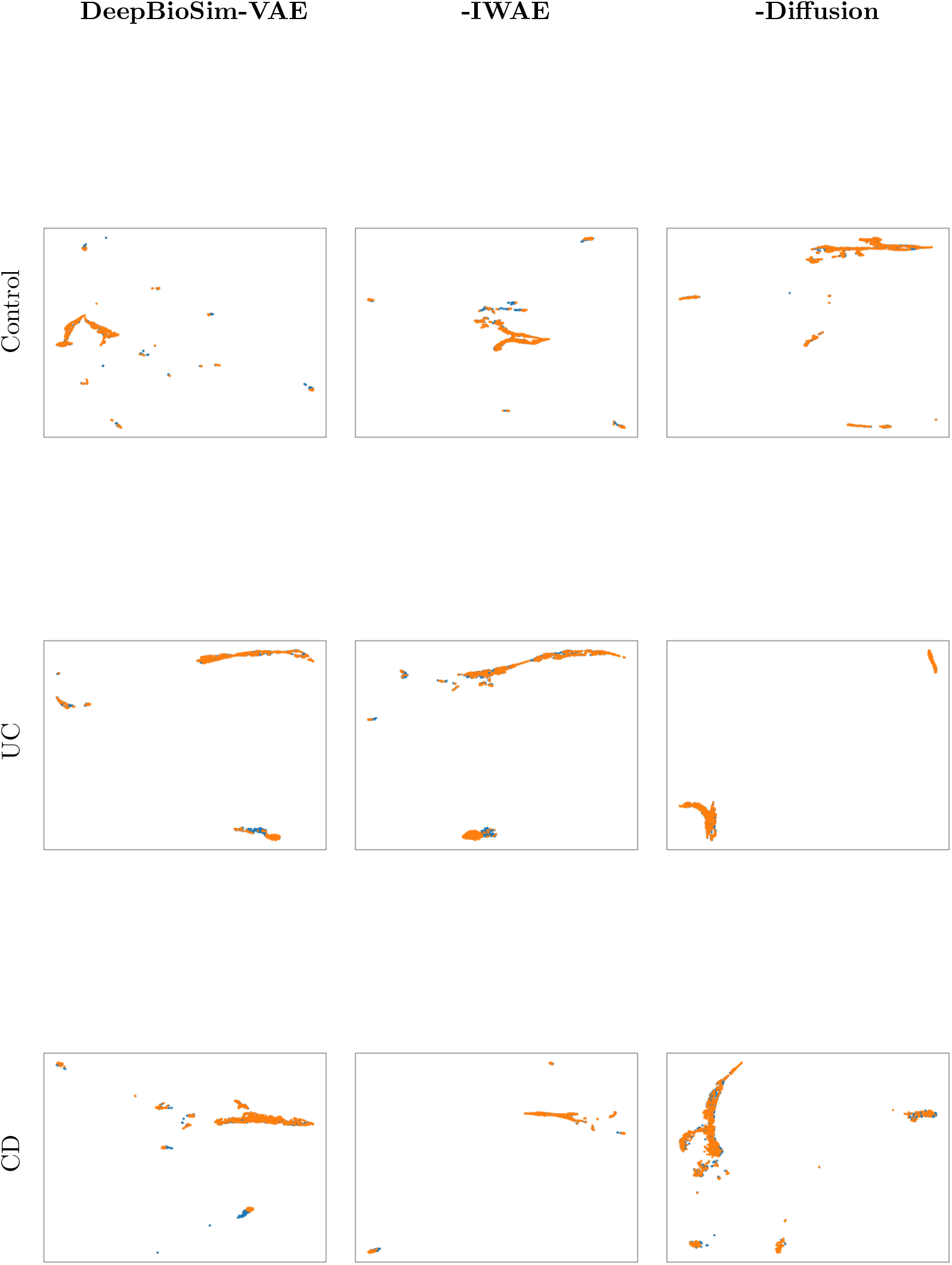
Gene-level concordance between the original human RNA-seq datasets from GSE165512 with different conditions and their three simulated counterparts, using UMAP visualization. Blue points represent genes in the original data; orange points represent genes in the simulated data.

## Appendix B Additional Sample Level Properties Evaluation

### B.1 Comparison of Sample-Level Properties Between the MOMS-PI 16S Dataset and Their Simulated Counterparts

**Fig. B9.**
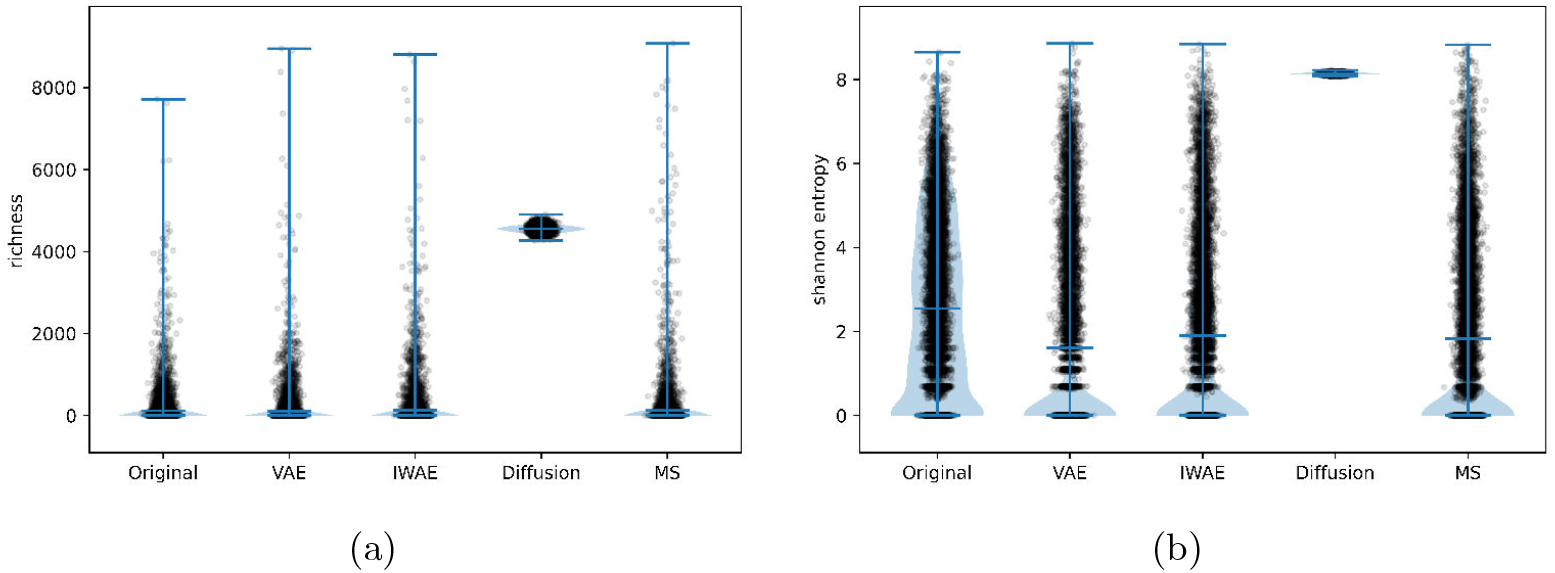
Sample-level alpha-diversity concordance between the original MOMS-PI 16S dataset and four simulated datasets. Panels (a) and (b) plot richness and Shannon entropy, respectively. Each point represents a single sample. MS refers to method MIDASim.

### B.2 Comparison of Sample-Level Properties Between the IBD 16S Dataset and Their Simulated Counterparts

**Fig. B10.**
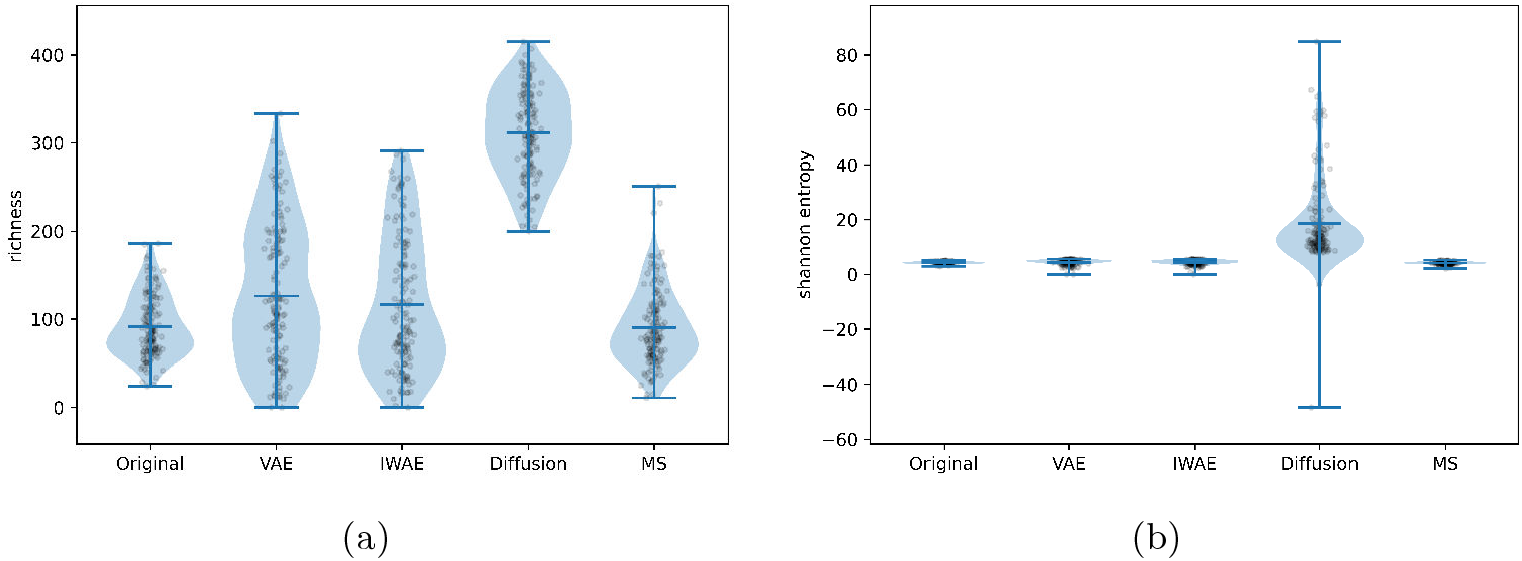
Sample-level alpha-diversity concordance between the original IBD 16S dataset and four simulated datasets. Panels (a) and (b) plot richness and Shannon entropy, respectively. Each dot represents a single sample.

### B.3 Comparison of Sample-Level Properties Between the Original Belstrom et al. Datasets and Their Simulated Counterparts

**Fig. B11.**
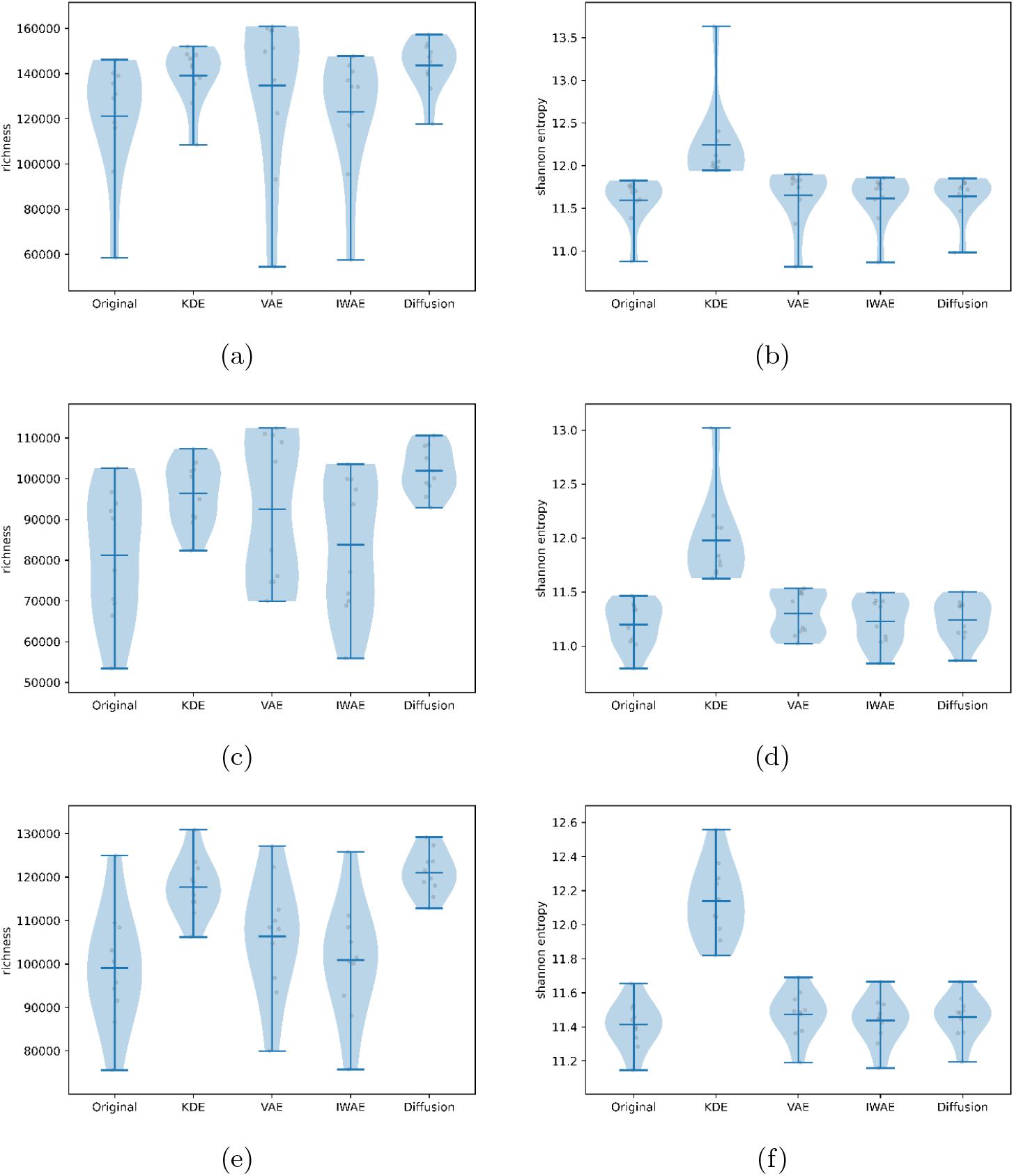
Sample-level alpha-diversity concordance between the original Belstrom datasets and two simulations. Panels a and b plot richness and Shannon entropy, respectively, for BelstromHealthy MG. Panels c and d repeat the richness and Shannon-entropy comparisons for BelstromCaries MG. Panels e and f repeat the richness and Shannon-entropy comparisons for BelstromPeriodontitis MG. Each dot represents a single sample.

**Fig. B12.**
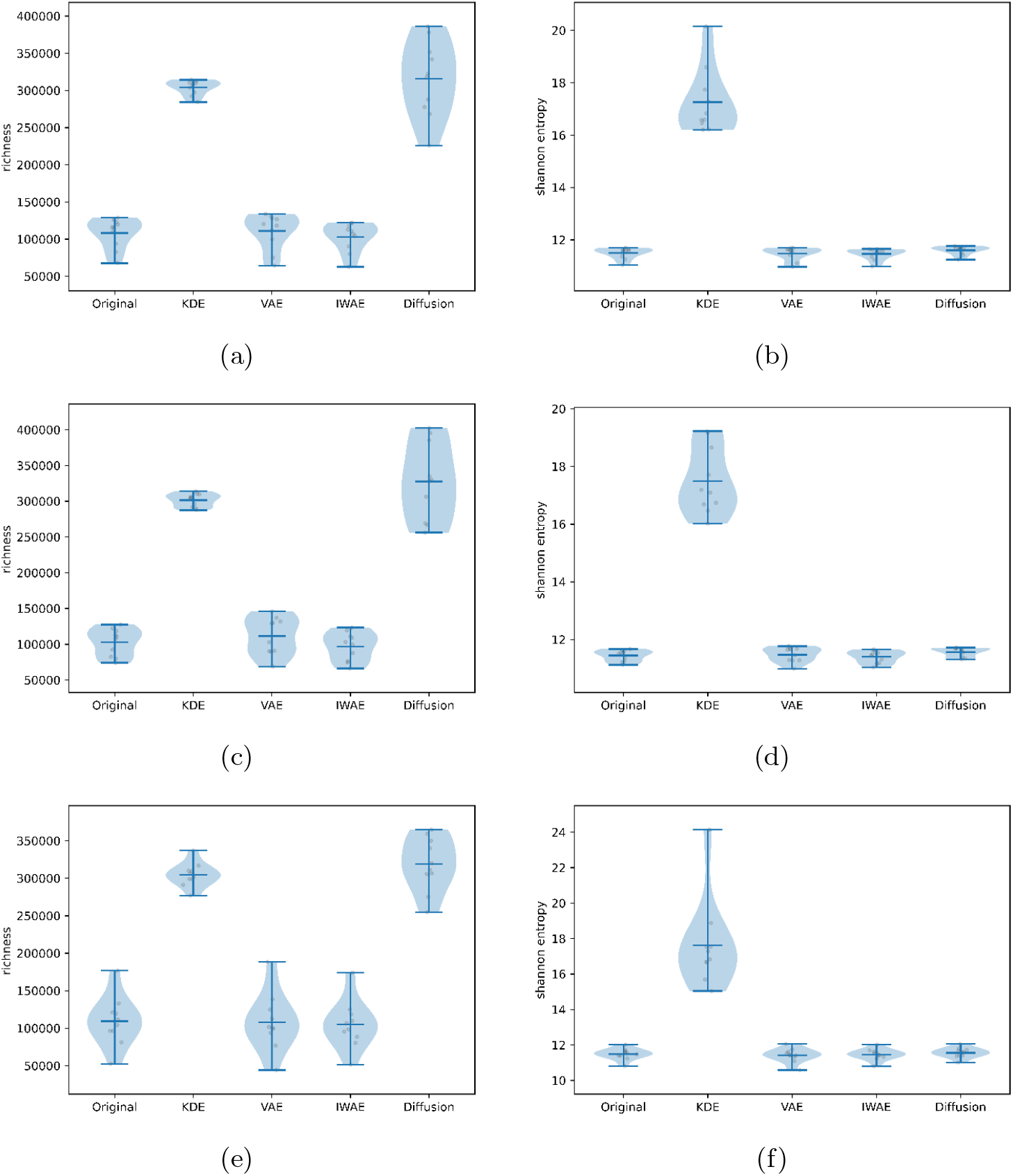
Sample-level alpha-diversity concordance between the original Belstrom datasets and two simulations. Panels a and b repeat the richness and Shannon-entropy comparisons for BelstromHealthy MT. Panels c and d repeat the richness and Shannon-entropy comparisons for BelstromCaries MT. Panels e and f repeat the richness and Shannon-entropy comparisons for BelstromPeriodontitis MT. Each dot represents a single sample.

**Table B1.**
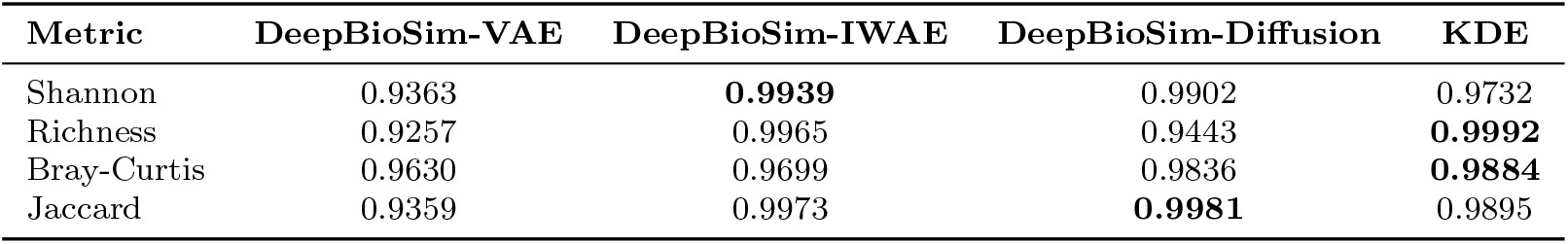
Pearson Correlation Coefficients of Diversity Index between Simulated BelstromHealthy-MG Dataset and Original Dataset

**Table B2.** Pearson Correlation Coefficients of Diversity Index between Simulated BelstromCaries-MG Datasets and Original Dataset

| Metric | DeepBioSim-VAE | DeepBioSim-IWAE | DeepBioSim-Diffusion | KDE |
| --- | --- | --- | --- | --- |
| Shannon | 0.9527 | **0.9734** | 0.9064 | 0.9534 |
| Richness | 0.9250 | 0.9914 | 0.7915 | **0.9993** |
| Bray-Curtis | 0.9780 | 0.9919 | **0.9926** | 0.9910 |
| Jaccard | 0.8970 | 0.9906 | **0.9914** | 0.9827 |

**Table B3.** Pearson Correlation Coefficients of Diversity Index between Simulated BelstromPeriodontitis-MG Datasets and Original Dataset

| Metric | DeepBioSim-VAE | DeepBioSim-IWAE | DeepBioSim-Diffusion | KDE |
| --- | --- | --- | --- | --- |
| Shannon | **0.9770** | 0.5994 | 0.9488 | 0.9706 |
| Richness | 0.9284 | 0.9879 | 0.7046 | **0.9976** |
| Bray-Curtis | 0.9720 | 0.9760 | 0.9807 | **0.9892** |
| Jaccard | 0.9471 | 0.9921 | **0.9920** | 0.9780 |

**Table B4.** Pearson Correlation Coefficients of Diversity Index between Simulated BelstromHealthy-MT Datasets and Original Dataset

| Metric | DeepBioSim-VAE | DeepBioSim-IWAE | DeepBioSim-Diffusion | KDE |
| --- | --- | --- | --- | --- |
| Shannon | 0.8809 | 0.1968 | -0.0462 | **0.9230** |
| Richness | 0.9857 | 0.9967 | 0.3979 | **0.9988** |
| Bray-Curtis | 0.9240 | -0.2857 | 0.8304 | **0.9507** |
| Jaccard | 0.9667 | 0.9916 | **0.9924** | 0.8473 |

**Table B5.** Pearson Correlation Coefficients of Diversity Index between Simulated BelstromCaries-MT Datasets and Original Dataset

| Metric | DeepBioSim-VAE | DeepBioSim-IWAE | DeepBioSim-Diffusion | KDE |
| --- | --- | --- | --- | --- |
| Shannon | 0.7602 | -0.0072 | -0.4061 | **0.8305** |
| Richness | 0.9486 | 0.9962 | -0.6382 | **0.9992** |
| Bray-Curtis | 0.9339 | 0.6072 | 0.9000 | **0.9766** |
| Jaccard | 0.9478 | **0.9867** | 0.9805 | 0.6955 |

**Table B6.** Pearson Correlation Coefficients of Diversity Index between Simulated BelstromPeriodontitis-MT Datasets and Original Dataset

| Metric | DeepBioSim-VAE | DeepBioSim-IWAE | DeepBioSim-Diffusion | KDE |
| --- | --- | --- | --- | --- |
| Shannon | **0.9333** | 0.8215 | 0.5614 | 0.7671 |
| Richness | 0.9764 | 0.9934 | 0.6439 | **0.9996** |
| Bray-Curtis | 0.9188 | 0.9548 | 0.8741 | **0.9681** |
| Jaccard | 0.9139 | 0.9930 | **0.9945** | 0.7468 |

### B.4 Sample-level properties comparisons of the original TCGA-HNSC RNA-seq data and simulated counterpart

**Fig. B13.**
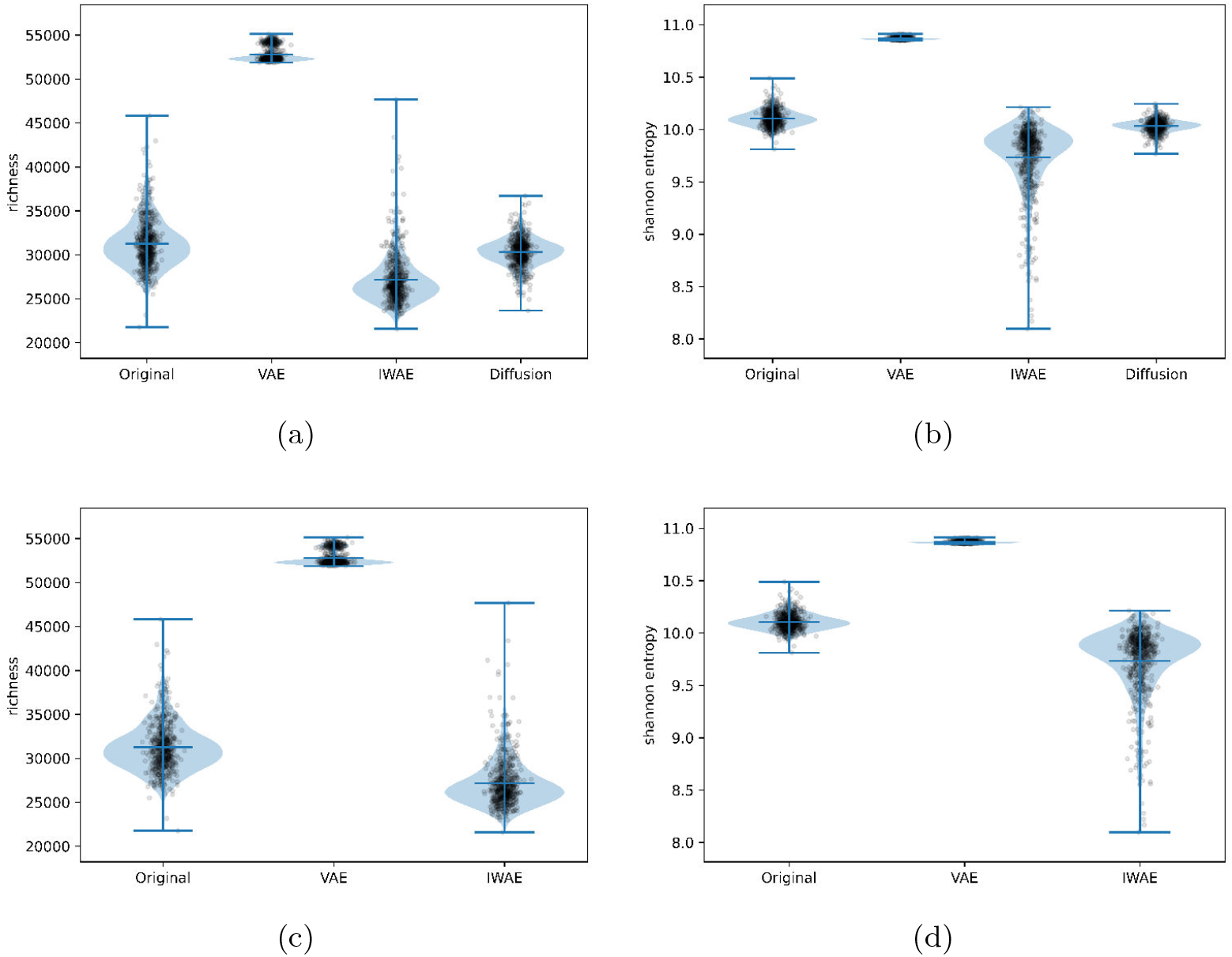
Sample-level alpha-diversity concordance between the original TCGA-HNSC RNA-seq dataset and three simulation methods. Panels a and b plot richness and Shannon entropy, respectively. Panels c and d repeat the richness and Shannon-entropy comparisons after omitting the DeepBioSim-Diffusion model to improve visual clarity. Each dot represents a single sample.

**Table B7.** Pearson Correlation Coefficients between Simulated TCGA-HNSC RNA-seq Datasets and Original Dataset

| Metric | DeepBioSim-VAE | DeepBioSim-IWAE |
| --- | --- | --- |
| Shannon | **0.3149** | 0.0768 |
| Richness | **0.9729** | 0.9644 |
| Bray-Curtis | **0.7316** | 0.6221 |
| Jaccard | 0.8483 | **0.8579** |

### B.5 Sample-level properties comparisons of the original dataset of GSE165512 and simulated Counterparts

**Fig. B14.**
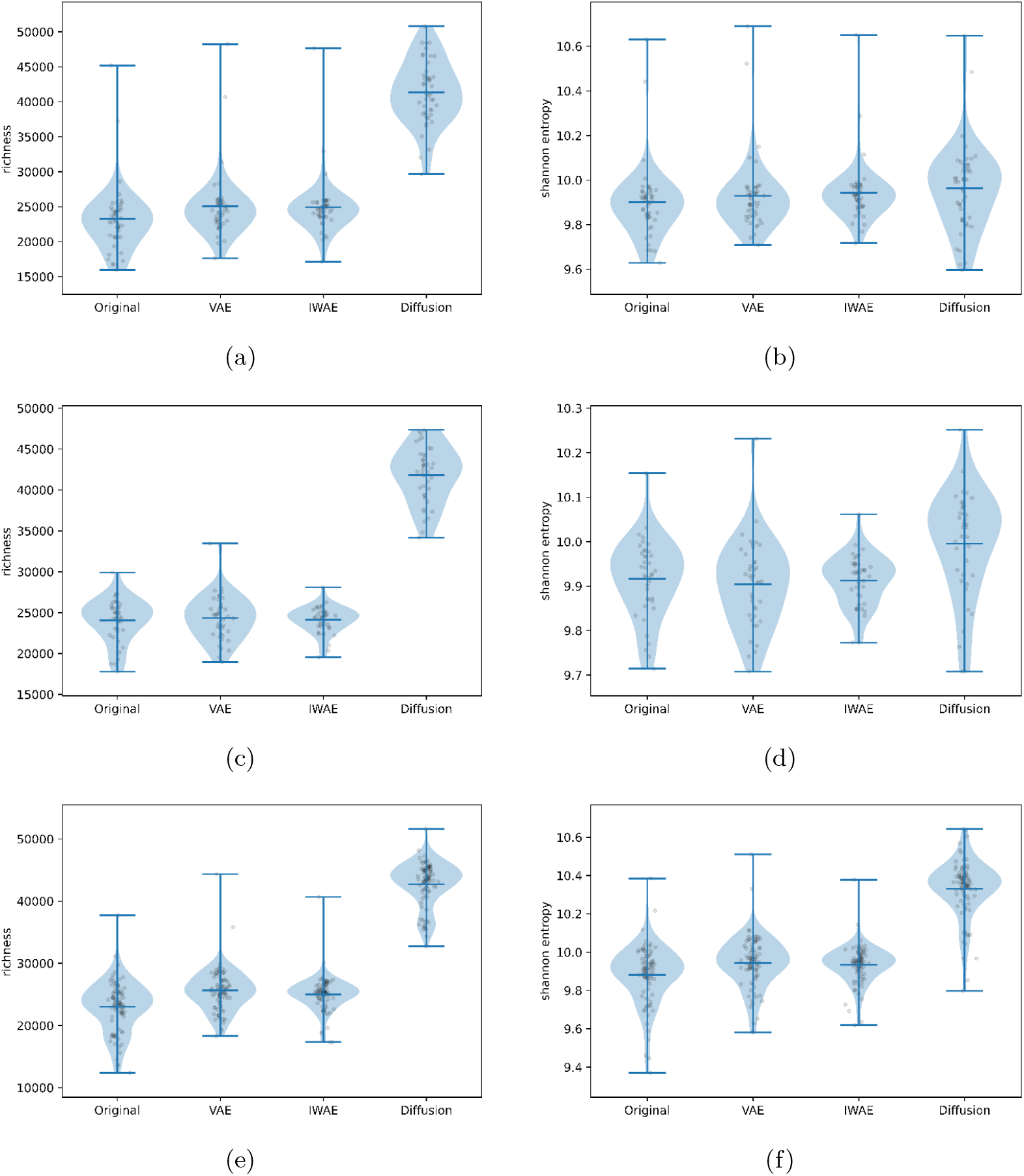
Sample-level alpha-diversity concordance between the original MOMS-PI 16S dataset and four simulation methods. Panels a and b plot richness and Shannon entropy, respectively, for control samples. Panels c and d repeat the richness and Shannon-entropy comparisons for UC samples. Panels e and f repeat the richness and Shannon-entropy comparisons for CD samples. Each dot represents a single sample.

**Table B8.** Pearson Correlation Correlation Coefficients of the Diversity Index between Simulated GSE165512-Control-RNA-seq Datasets and Original Dataset

| Metric | DeepBioSim-VAE | DeepBioSim-IWAE | DeepBioSim-Diffusion |
| --- | --- | --- | --- |
| Shannon | **0.8248** | -0.3351 | 0.6416 |
| Richness | **0.9212** | 0.9174 | 0.6182 |
| Bray-Curtis | **0.8669** | 0.5573 | 0.4281 |
| Jaccard | 0.9541 | 0.9559 | **0.9643** |

**Table B9.** Pearson Correlation Correlation Coefficients of the Diversity Index between Simulated GSE165512-UC-RNA-seq Datasets and Original Dataset

| Metric | DeepBioSim-VAE | DeepBioSim-IWAE | DeepBioSim-Diffusion |
| --- | --- | --- | --- |
| Shannon | 0.5861 | **0.7474** | 0.2206 |
| Richness | 0.8985 | **0.9357** | 0.7542 |
| Bray-Curtis | 0.5106 | **0.8122** | 0.3479 |
| Jaccard | 0.9129 | 0.9495 | **0.9558** |

**Table B10.** Pearson Correlation Coefficients of the Diversity Index between Simulated GSE165512-CD-RNA-seq Datasets and Original Dataset

| Metric | DeepBioSim-VAE | DeepBioSim-IWAE | DeepBioSim-Diffusion |
| --- | --- | --- | --- |
| Shannon | **0.8248** | -0.3351 | 0.6416 |
| Richness | **0.9212** | 0.9174 | 0.6182 |
| BC | **0.8669** | 0.5573 | 0.4281 |
| Jaccard | 0.9541 | 0.9559 | **0.9643** |

